# Light-harvesting by antenna-containing rhodopsins in pelagic Asgard archaea

**DOI:** 10.1101/2024.09.18.613612

**Authors:** Gali Tzlil, María del Carmen Marín, Yuma Matsuzaki, Probal Nag, Shota Itakura, Yosuke Mizuno, Shunya Murakoshi, Tatsuki Tanaka, Shirley Larom, Masae Konno, Rei Abe-Yoshizumi, Ana Molina-Márquez, Daniela Bárcenas-Pérez, José Cheel, Michal Koblížek, Rosa León, Kota Katayama, Hideki Kandori, Igor Schapiro, Wataru Shihoya, Osamu Nureki, Keiichi Inoue, Andrey Rozenberg, Ariel Chazan, Oded Béjà

## Abstract

Aquatic bacterial rhodopsin proton pumps have been recently reported to utilize hydroxylated carotenoids^1,2^. Here, by combining a marine chromophore extract with purified archaeal rhodopsins identified in marine metagenomes, we report on light energy transfer from diverse hydroxylated carotenoids (lutein, diatoxanthin, and fucoxanthin) to heimdallarchaeial rhodopsins (HeimdallRs)^3,4^ from uncultured marine planktonic members of the “*Ca.* Kariarchaeaceae” (“*Ca.* Asgardarchaeota”)^5^. These light-harvesting antennas absorb in the blue-light range and transfer energy to the green-light absorbing retinal chromophore within HeimdallRs. Furthermore, antenna enhancement of proton pumping by HeimdallRs is also observed under white-light illumination along with a carotenoid-binding–induced structural change in the protein. Our results indicate that the use of light-harvesting antennas in microbial rhodopsins is observed not only in bacteria but also in marine archaea.

---

Proteorhodopsins (PRs) and xanthorhodopsins (XRs) are microbial light-driven proton pumps identified in widespread and abundant bacterial, archaeal and some eukaryotic groups, in marine and freshwater environments^6^. It has been predicted that about 50% of microbes in the ocean’s photic zone possess these rhodopsins^7^ and use them to harvest light energy to perform phototrophy^8,9^.

The phenomenon of energy transfer from ketolated carotenoids to retinal in some XRs was first demonstrated two decades ago^10,11^. Recently, based on functional metagenomics combined with chromophore extraction from the environment, it was estimated that in the oceans about a third of PRs and a fifth of XRs can potentially use carotenoids as light-harvesting antennas^2^; suggesting a substantial impact on rhodopsin-mediated phototrophy in marine environments. These antennas transfer energy in the violet- and blue-light range (∼400-500 nm) to the green-absorbing retinal chromophore (∼550 nm) within these rhodopsins^2,10,11^, therefore enabling them to use light that is otherwise not available to them. The energy transfer is facilitated by a lateral opening in the rhodopsin (known as the fenestration), exposing the retinal β-ionone ring to one of the rings of the cyclic carotenoid antenna^2,11,12^, with rhodopsins lacking this fenestration failing to bind carotenoids^2,11^.

Thus far, direct light energy transfer from hydroxylated carotenoids (lutein, zeaxanthin and nostoxanthin) has been reported only for bacterial PRs and XRs^1,2^. In this study, we searched for rhodopsin-carotenoid complexes originating from marine archaea. Several marine group II and III archaea (MGII/III, “*Ca.* Poseidoniia”)^13^ contain both PRs and light-driven proton pumps ACB (*Archaea* Clade B) rhodopsins^14–17^ (Fig. 1a, see light-driven proton pumping activity for ACB-G35 rhodopsin in Supplementary Fig. 1). While PRs from “*Ca.* Poseidoniia” have bulky residues at the fenestration-enabling position G156 in transmembrane helix 5 (TM5) and are not fenestrated, most ACB rhodopsins do possess a fenestration in some proximity to the retinal ring facilitated by the presence of the unusual for this position leucine residue (Fig. 1a,b). In addition to rhodopsins from “*Ca.* Poseidoniia”, a proton-pumping rhodopsin was recently reported for “*Ca.* Heimdallarchaeia” (“*Ca.* Asgardarchaeota”)^3,4^. This protein (referred here as HeimdallR1) has glycine at position G156 and is predicted to have a fenestration exposing the retinal ring similarly to *Salinibacter* XR^12^ and Kin4B8-XR^2^. These observations hint at the possibility that some of these strictly archaeal proton pumps might be able to utilize carotenoid antennas, despite the evolutionary distance separating them from the more common and predominantly bacterial PR and XR families and the presence of non-canonical motifs in transmembrane helix 3 (TM3) (see Fig. 1a). The appearance of proton-pumping rhodopsins with a potential for antenna binding in “*Ca.* Heimdallarchaeia” is of particular interest as members of this Asgard group are the closest archaeal relatives of eukaryotes ^18,19^ and little is known about photoheterotrophy among them.

**Fig. 1.**
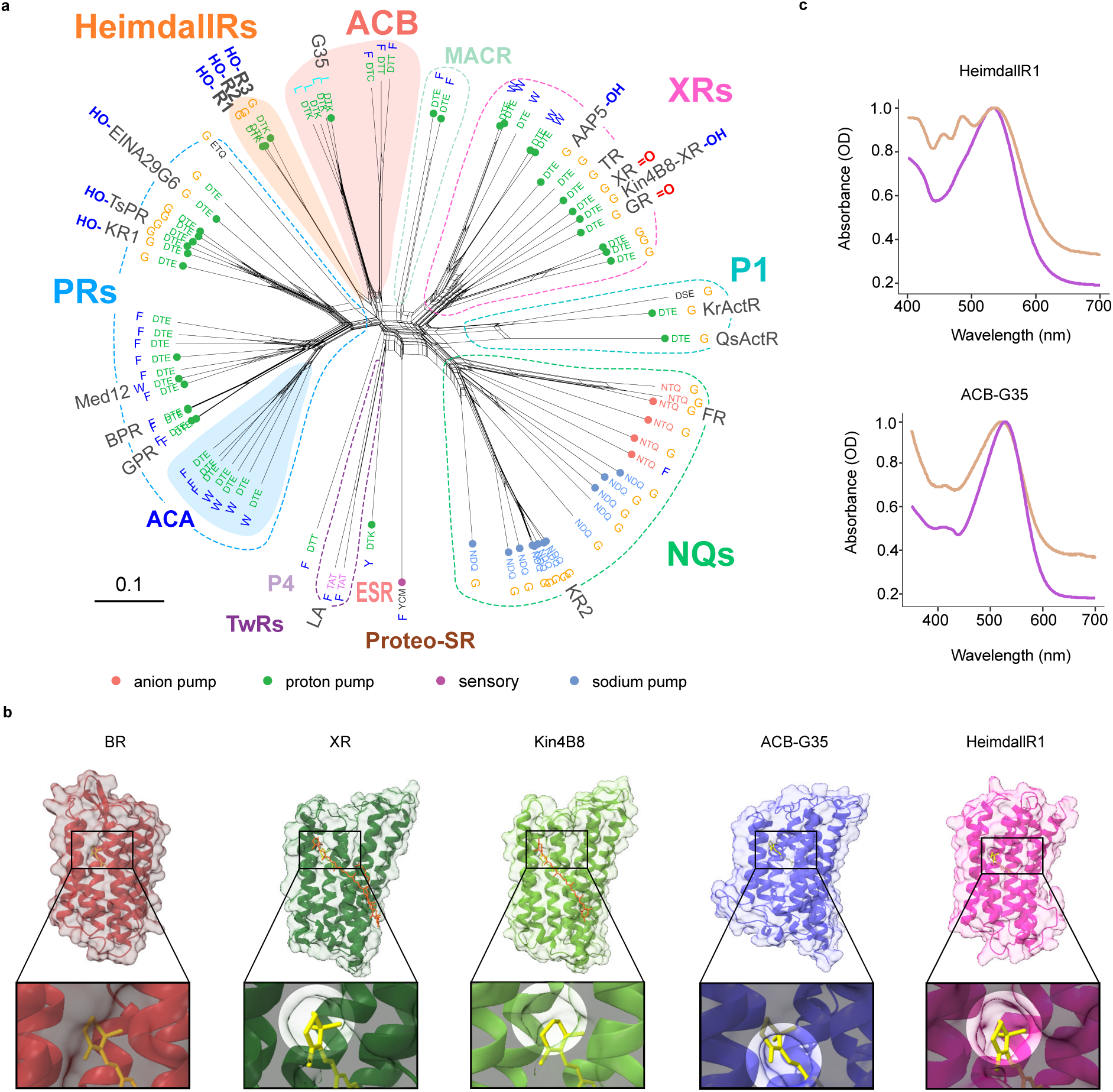
Evolutionary ties and xanthophyll binding potential in fenestrated archaeal rhodopsin proton pumps. **a**, NeighborNet network of PRs, XRs, ACB rhodopsins, HeimdallRs and related rhodopsin families. The three clades appearing in marine archaea are highlighted: *Archaea* clade A PRs (ACA) among marine group II and marine group III (*Thermoplasmatota*: “*Ca.* Poseidoniia”), the *Archaea* clade B family (ACB) among marine group II and the family of HeimdallRs among the “*Ca.* Kariarchaeaceae” (“*Ca.* Asgardarchaeota”: “*Ca.* Heimdallarchaeia”). Indicated are (from inside out): activity, TM3 motif, residue at the fenestration position G156 in TM5, alias and type of carotenoid antenna (-OH for xanthophylls with hydroxyl at carbon C3 and =O for xanthophylls with keto-group at carbon C4). **b**, Structural comparison of fenestrated rhodopsins: *S. ruber* XR (PDB: 3DDL)^12^, Kin4B8-XR (PDB: 7YTB)^2^, ACB-G35 rhodopsin (AlphaFold2 model)^20^ and HeimdallR1 (AlphaFold2 model) versus a non-fenestrated rhodopsin — bacteriorhodopsin (BR; PDB: 1IW6)^21^. The structures are aligned based on the retinal β-ionone ring position, and the fenestration zone is highlighted. For clarity, salinixanthin and zeaxanthin are not shown in the enlarged fenestration zone of *S. ruber* XR and Kin4B8-XR structures, respectively. **c**, Absorbance spectra of ACB-G35 rhodopsin and HeimdallR1 before (purple) and after (brown) incubation, and wash, with a marine chromophore extract.

For the first exploratory experiment, two representative fenestrated archaeal rhodopsins, ACB-G35 rhodopsin and HeimdallR1 (Fig. 1b) were incubated with a marine chromophore extract containing a natural mix of carotenoids. After repurifying the proteins from the chromophore extract mixture, a change in the absorbance spectrum was observed for HeimdallR1 but not for the ACB rhodopsin (Fig. 1c), suggesting binding of specific chromophores to HeimdallR1. High-performance liquid chromatography with diode array detector (HPLC-DAD) analysis of the complexes showed that the enriched chromophores consisted mainly of cyclic hydroxylated (lutein, diatoxanthin, and fucoxanthin) and non-hydroxylated (β-carotene) carotenoids (see Extended Data Fig. 1; see carotenoid structures in Fig. 2a). When switching to pure carotenoids, the nonsymmetric hydroxylated carotenoids lutein, diatoxanthin and fucoxanthin showed binding to HeimdallR1, while β-carotene did not (Fig. 2a). Interestingly, this carotenoid specificity of HeimdallR1 is similar but not identical to that of the bacterial Kin4B8-XR which does not bind fucoxanthin (Extended Data Fig. 2). This indicates a unique binding preference of HeimdallR1 toward fucoxanthin and further implies that carotenoid binding in fenestrated rhodopsins is a complex trait.

**Fig. 2.**
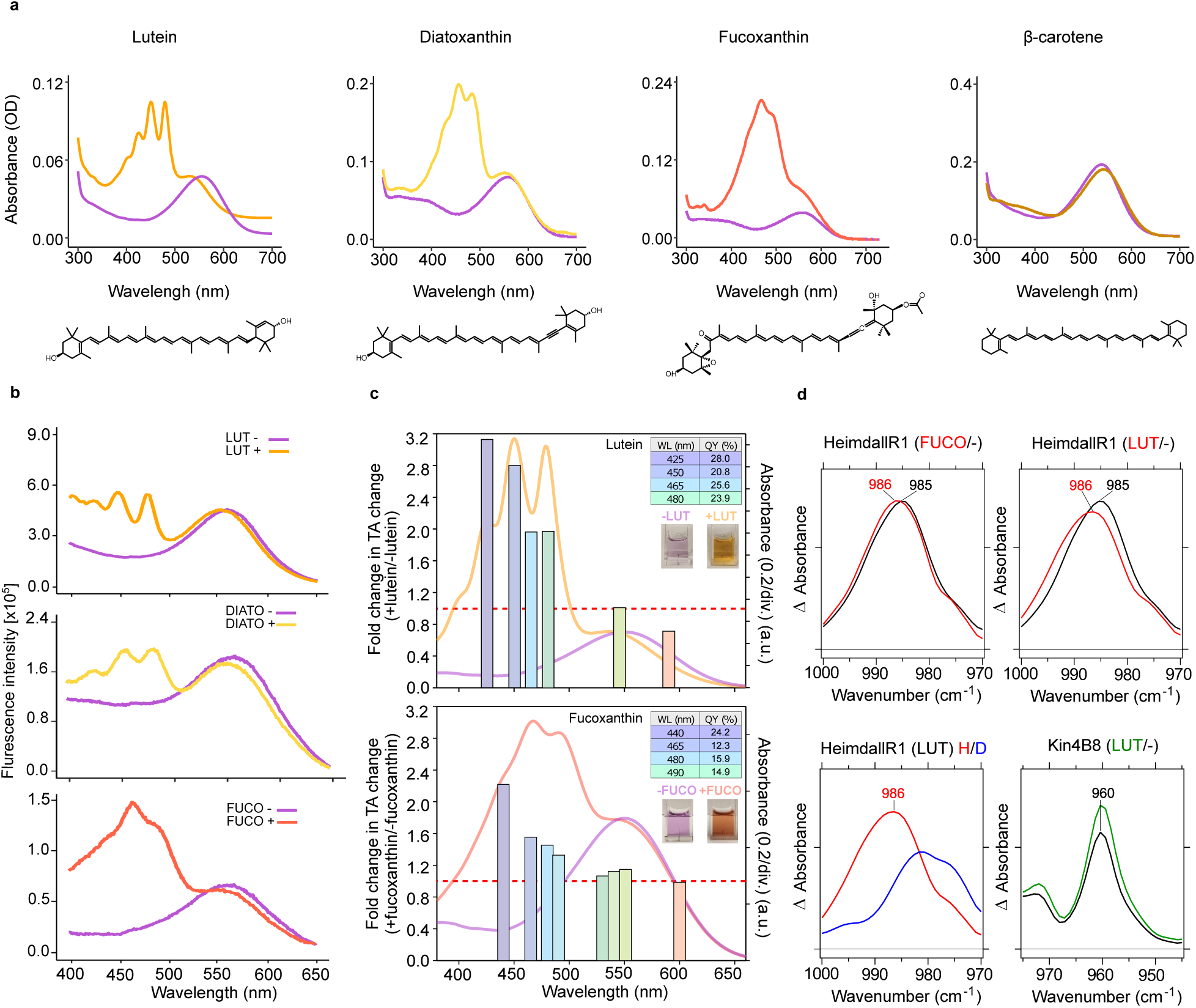
Spectroscopic characterics of HeimdallR1 bound to different xanthophylls. **a**, Absorbance changes of HeimdallR1 before (purple) and after (orange) incubation, and wash, with pure lutein, diatoxanthin, fucoxanthin and β-carotene. Carotenoid structures are indicated below. **b**, Fluorescence excitation spectra of HeimdallR1 upon incubation with (orange) or without (purple) for lutein (top), diatoxanthin (middle), or fucoxanthin (bottom); emission was recorded at 720 nm. **c**, The ratios of transient absorption (TA) change in HeimdallR1 with and without lutein (top) and with and without fucoxanthin (bottom) at different excitation wavelengths (425, 450, 465, 480, 545, and 590 nm for lutein and 440, 465, 580, 490, 530, 540, 550, and 600 nm for fucoxanthin) (bars colored according to the color of excitation light). The absorption spectra of HeimdallR1 without (purple line) and with (orange line for lutein and coral for fucoxanthin) xanthophylls were overlaid. The red dashed line indicates no difference between with and without xanthophyll. Tables with the QY percentages and the pictures of the purified proteins are shown next to the corresponding results. **d**, Light-minus-dark difference FTIR spectra at 77 K upon illumination of HeimdallR1 with (red) or without (black) fucoxanthin, HeimdallR1 with (red) or without (black) lutein, HeimdallR1 with lutein in H_2_O (red) or D_2_O (blue), and Kin4B8-XR with (green) or without (black) lutein. Hydrated films of lipid-reconstituted protein with H_2_O are illuminated at 540 nm light (solid lines), which forms the red-shifted K intermediate. Each peak originates from a hydrogen out-of-plane (HOOP) vibration of the retinal chromophore, which shifts upon xanthophyll binding to HeimdallR1, but not to Kin4B8-XR. One division of the y-axis corresponds to 0.0006 absorbance units. Lutein (LUT), diatoxanthin (DIATO), fucoxanthin (FUCO).

Further experiments demonstrated that the carotenoid binding in HeimdallR1 is coupled to energy transfer. Energy transfer from lutein or fucoxanthin to the retinal molecule of HeimdallR1 was indicated by fluorescence measurements (Fig. 2b). Analogously, laser-flash photolysis revealed an enhancement in transient absorption signal for HeimdallR1 bound to lutein or fucoxanthin when excited at the violet-blue (425-490 nm) region, enhancing the retinal isomerization quantum yield (Fig. 2c). Moreover, HeimdallR1 photocycle was enhanced to a greater extent (by 3.1 folds with lutein and 2.2 with fucoxanthin) when bound to xanthophylls, compared to the photocycle of Kin4B8-XR bound to lutein (1.6 folds enhancement)^2^.

The influence of the xanthophyll binding on the retinal isomer composition in HeimdallR1 was investigated using HPLC of retinal oximes produced by hydrolyzing the retinal Schiff base (RSB) with hydroxylamine. Most of the retinal chromophores in the dark-adapted (DA) protein showed an all-*trans* configuration (Extended Data Fig 3a), and the binding of lutein and fucoxanthin further increased the fraction of the all-*trans*-retinal form. The retinal photoisomerized to the 13-*cis* form, which was reflected in transient absorption changes, representing red-shifted (K and O) and blue-shifted (M) photointermediates (Extended Data Fig. 3b, c, and d) in addition to L, N, and HeimdallR1ʹ intermediates with maximum absorption wavelengths (λ^a^_max_) close to the initial state (Extended Data Fig. 3e). A sharp peak at 471 and 480 nm was observed when lutein or fucoxanthin, respectively, were bound to the protein (asterisks in Extended Data Fig. 3c and e), indicating that a large conformational change of the protein occurs during the photocycle, which likely alters xanthophyll structures (Extended Data Fig. 3f).

To investigate how the retinal binding pocket is affected by xanthophyll binding, we purified HeimdallR1 without carotenoids, as well as with lutein or fucoxanthin (Extended Data Fig. 4a). Binding of both xanthophylls elicited a blue-shift in the λ^a^_max_ compared to HeimdallR1 alone, which was not observed with the bacterial rhodopsin proton pumps^2,11^. Meanwhile, a small λ^a^_max_ shift was observed on the acidic side of HeimdallR1 alone, whereas HeimdallR1 with lutein or with fucoxanthin exhibited larger red shifts (20 nm and 11 nm, respectively, Extended Data Fig. 4b). This effect is caused by the protonation of the counterion in the third transmembrane helix (TM3) as known for many microbial rhodopsins. The retinal counterion of HeimdallR1, D81, is fully protonated at pH < 4.0 irrespective of xanthophyll binding. Hence, the difference in the λ^a^_max_ shift is due to the difference in the degree of counterion protonation at neutral pH, and D81 of HeimdallR1 binding lutein is more deprotonated than that of HeimdallR1 alone at pH 7.0 (Extended Data Fig. 4c). Since the deprotonated counterion can act as the proton acceptor to receive H^+^ from the RSB^22^, the higher deprotonation suggests that lutein binding would enhance proton pumping, which aligns with the large M formation observed for HeimdallR1 binding lutein (Extended Data Fig. 3e).

To gain further structural insights into the xanthophyll binding, we measured light-induced differences in Fourier transform infrared (FTIR) spectra of HeimdallR1 with or without lutein or fucoxanthin at 77 K. Extended Data Fig. 5a and b show UV-visible spectra and formation of the K intermediate at 77 K, respectively. While light-induced difference FTIR spectra looked similar for HeimdallR1 with or without xanthophylls (Extended Data Fig. 5c), we did observe two xanthophyll-dependent frequency shifts. One was seen in the C=C stretching frequency of the retinal chromophore (Extended Data Fig. 5d), which is however not due to structural reasons, as C=C stretch reflects λ^a^_max_ (Extended Data Fig. 5b). Indeed, the spectra of the C-C stretch region (Extended Data Fig. 5e) and amide-I (Extended Data Fig. 5h) were coincident, indicating that xanthophyll binding has no effect on chromophore structure and peptide backbone, respectively. The second shift was observed for a hydrogen out-of-plane (HOOP) band of the retinal chromophore in the K intermediate of HeimdallR1, while no such shift was observed for Kin4B8-XR (Fig. 2d and Extended Data Fig. 5f). As HOOP bands appear by chromophore distortions, we conclude that binding of lutein or fucoxanthin to HeimdallR1 influences the retinal distortion in the K intermediate. As the HOOP band is downshifted by an H/D exchange (Fig. 2d and Extended Data Fig. 5g), the chromophore distortion is located near the Schiff base. This means that xanthophyll binding to HeimdallR1 alters the structure of the retinal near the Schiff base region, in other words, on the side of the retinal moiety furthest from the fenestration. This is in clear contrast to Kin4B8-XR, which binds xanthophylls without affecting structural changes upon retinal photoisomerization^2^.

The influence of carotenoid binding on the efficiency of proton pumping of HeimdallR1 was investigated by expressing the protein in *E. coli* spheroplasts. When illuminated with white light (400-700 nm), the spheroplasts demonstrated an enhanced light-dependent outward proton flux upon the addition of lutein or fucoxanthin but not of diatoxanthin (Fig. 3a top). Introduction of a fenestration-blocking mutation, G141F (XR position Gly156), abolished carotenoid binding (Extended Data Fig. 2c). Under the same illumination conditions and in the presence of lutein, diatoxanthin or fucoxanthin, the pumping activity of the non-fenestrated HeimdallR-G141F was reduced significantly (Fig. 3a bottom). This decline is probably caused by non-specific adsorption of the carotenoids to the spheroplasts, resulting in masking of light. This implies that under our experimental settings, the observed proton flux in wild-type HeimdallR1 with all three xanthophyll antennas, under white light, is in effect underestimated. This would indicate that even when diatoxanthin serves as an antenna there is a significant enhancement in proton flux under these conditions. Importantly, the enhancement of the pumping activity in HeimdallR1 when bound to lutein was maintained over a range of decreasing light intensities (2,000, 860, 590 and 515 µmol m^−2^ s^-1^, Fig. 3b). This suggests that in the photic zone, under open ocean water conditions^23^, xanthophyll antennas have the potential to enhance rhodopsin performance from the very surface to tens of meters deep.

**Fig. 3.**
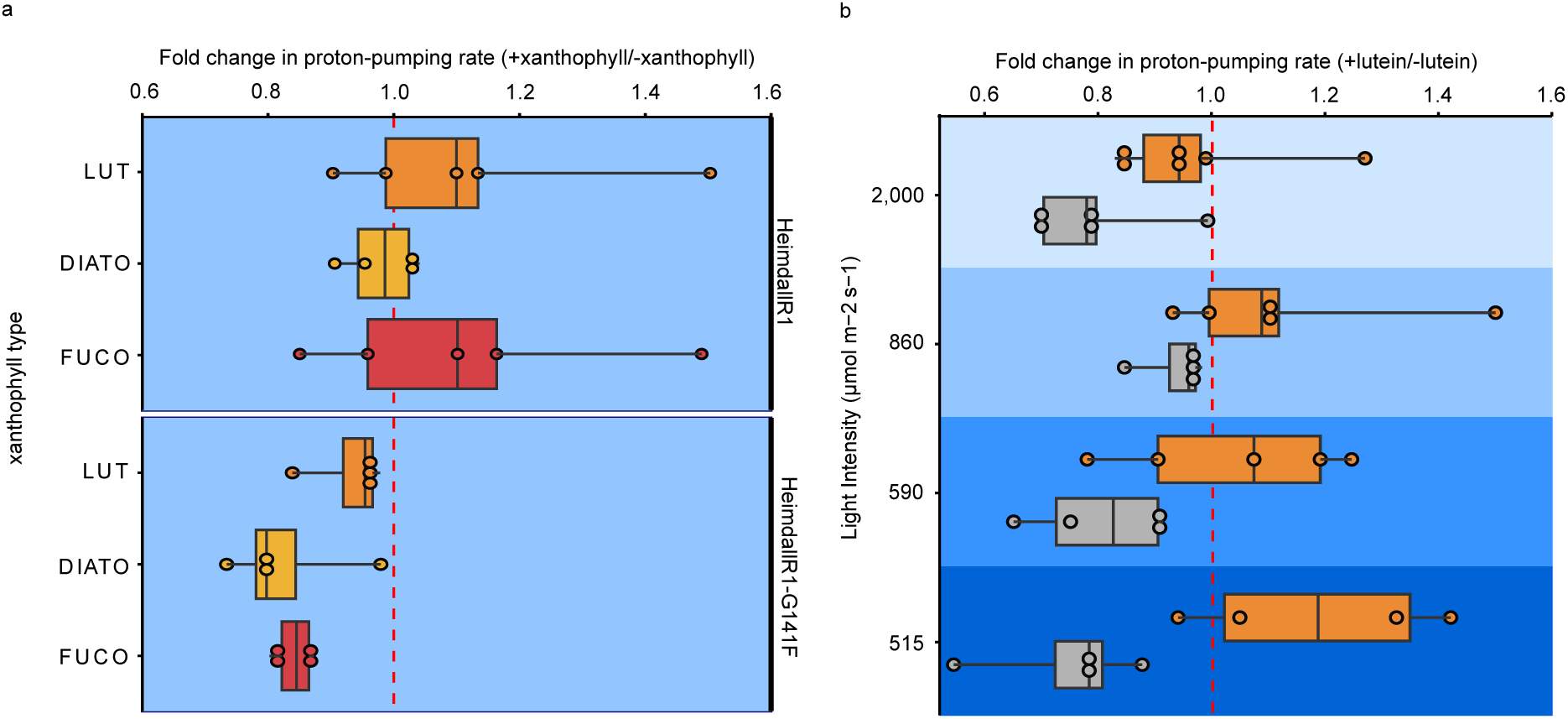
Proton pumping activity of HeimdallR1 under white-light illumination. **a**, Light-driven proton-pumping rates in HeimdallR1 and HeimdallR1-G141F *E. coli* spheroplasts with and without lutein, diatoxanthin or fucoxanthin (depicted in orange, mustard and red, respectively) under white-light illumination (400-700 nm, at 860 µmol m^−2^ s^−1^). Each dot represents the proton-pumping rate of HeimdallR1 and HeimdallR1-G141F with a xanthophyll over without the xanthophyll. Carotenoid names are abbreviated as follows: LUT – lutein, DIATO – diatoxanthin, FUCO – fucoxanthin. A total of 26 ratios were used. **b**, Light-driven proton-pumping rates in HeimdallR1 (depicted in orange circles) and HeimdallR1-G141F (depicted in gray circles) *E. coli* spheroplasts with and without lutein under various white-light intensities (400-700 nm, at 2,000, 860, 590 and 515 µmol m^−2^ s^−1^). Each dot represents the proton-pumping rate of HeimdallR1 and HeimdallR1-G141F with lutein over without lutein. A total of 37 ratios were used. In both panels, box plots represent the lower quartile, median and the upper quartile, and the whiskers depict 1.5 × the interquartile range. The red dashed lines correspond to no differences in proton-pumping activity relative to HeimdallR1 or HeimdallR1-G141F without a xanthophyll. Highly significant effects were found only for the factor of fenestration (wild type vs. G141F mutation) in both cases, see Supplementary Table 1.

To examine the unique xanthophyll binding preference observed for HeimdallRs, the structure of HeimdallR1 was analyzed. We succeeded in purifying the fucoxanthin-bound HeimdallR1 (Extended Data Fig. 6a), and determined the crystal structure of HeimdallR1 at a 2.0-Å resolution by the lipidic cubic phase method (Extended Data Fig. 6b, Supplementary Table 2). HeimdallR1 appeared as a monomer in the crystal packing (Extended Data Fig. 6c), and thus its physiological oligomeric structure remains unknown. The protein has the canonical architecture with seven transmembrane helices (TM1-7) and an all-*trans* retinal (ATR) (Fig. 4a,b, Extended Data Fig. 6d) and demonstrates an overall high similarity to PRs, XRs and related proteins. HeimdallR1 shows a peculiar deformation in TM6: a π-bulge followed by a 3_10_-helix segment, also known for GPR^24^ and ESR^25^ but absent in XRs, NQs and related families (Extended Data Fig. 6e), which makes it a prominent structural hallmark of PR-like families. HeimdallR1 diverges from the related families most conspicuously in the structure of the loop portion (Extended Data Fig. 6f,g). Notably, intracellular loop 3 (ICL3) forms a short membrane-extending α-helix and tightly interacts with TM6 (Fig. 4b, Extended Data Fig. 6h), which is not observed in other microbial rhodopsins. Another salient structural feature of HeimdallR1 is an unusually extensive cavity at the cytoplasmic side which directly connects the putative proton donor residue K92 with the bulk solvent which distinguishes it from ESR with which it shares the non-canonical cationic donor ^25,26^.

**Fig. 4.**
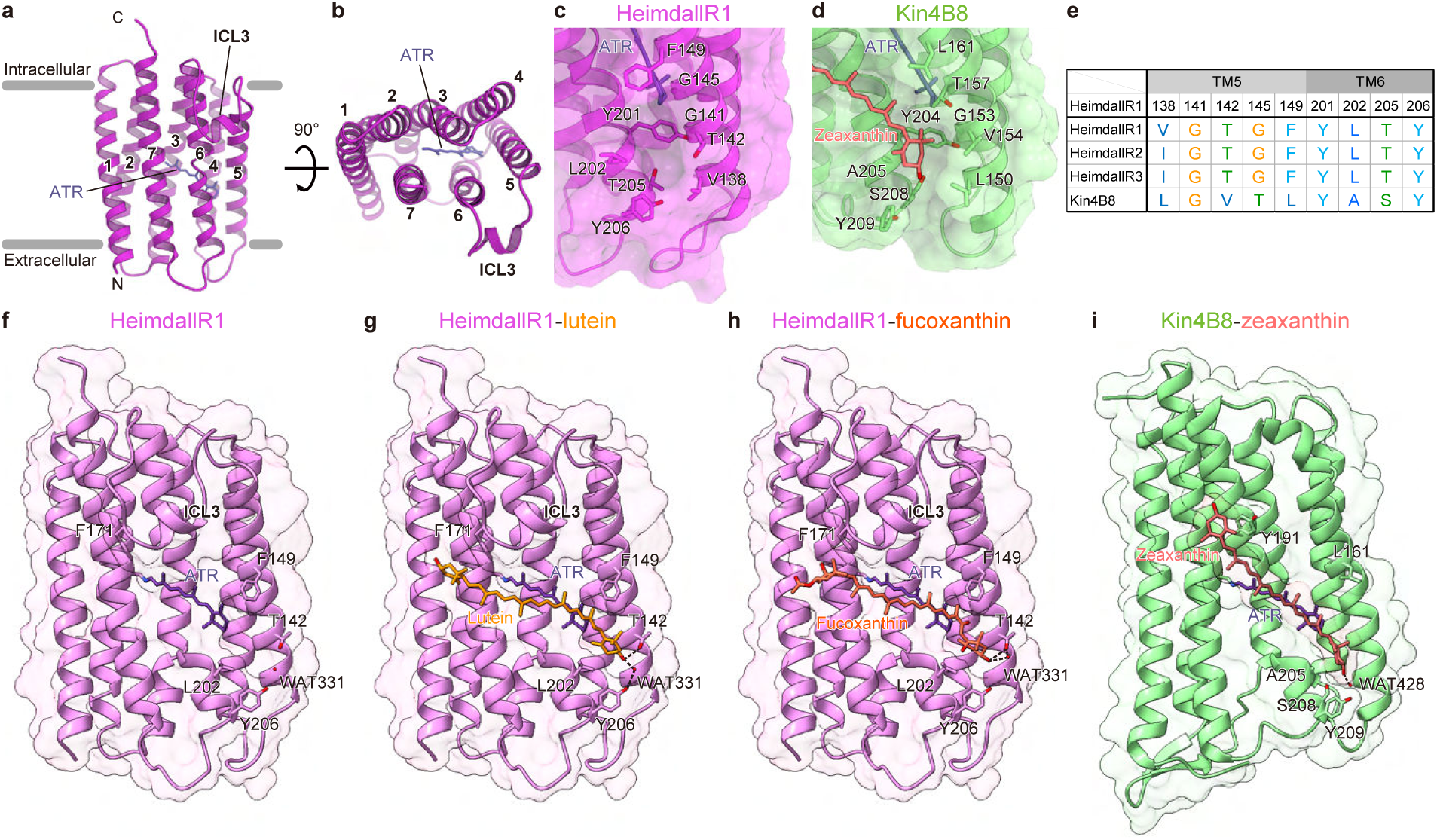
Crystal structure of HeimdallR1 and the docking of lutein and fucoxanthin using QM/MM simulations. **a**, **b**, Overall structure of HeimdallR1 with the retinal chromophore, viewed from the membrane plane (**a**) and the intracellular side (**b**). **c**, **d**, Fenestrations in HeimdallR1 (**c**) and Kin4B8-XR (PDB ID: 8I2Z) (**d**). **e**, Conservation of the residues surrounding the fenestrations in HeimdallRs and Kin4B8-XR. Glycine (G) and tyrosine (Y) are coloured orange and cyan, respectively, while polar [threonine (T), serine (S)] and hydrophobic [valine (V), isoleucine (I), leucine (L), alanine (A)] residues are coloured green and blue, respectively. **f**-**h**, HeimdallR1 structure, energy minimized using the hybrid QM/MM method (**f**), and the docking of lutein (**g**) and fucoxanthin (**h**) along the outer surface of TM6. Black dashed lines indicate hydrogen bonds. **i**, Zeaxanthin-bound Kin4B8 (PDB ID: 8I2Z). See Extended Data Fig. 7 for zoom-in on the fenestration area.

In line with the initial predictions, the structure of HeimdallR1 features a fenestration in the vicinity of the retinal owing to two glycine residues, G141 and G145 (Fig. 4c). The residues in the bottom of the fenestration are hydrophilic (Fig. 4d,e) as in Kin4B8-XR, thus favorable for carotenoid binding. However, the resolved crystal structure lacks the antenna indicating that fucoxanthin was dissociated under the conditions used for crystallization (Extended Data Fig. 6c). We thus explored the binding of xanthophylls (lutein and fucoxanthin) to HeimdallR1 using the hybrid quantum mechanics/molecular mechanics (QM/MM) method^27^.

The carotenoids can form an intricate network of hydrogen bonding with the water molecules present in the vicinity, and the polar residues present on TM5 and TM6 (T142 and Y206, respectively) via the hydroxyl groups present in the rings. Apart from this, the phenyl residue (F149 in TM5) can also stabilize the carotenoid via π-interaction. We observe that lutein can bind via both the rings, with the most favorable one being docking through the β-ionone ring (C5 atom of the ring directed towards retinal) where the hydroxyl group forms hydrogen bonds with WAT331 and T142, apart from the C-H…π interaction of a methyl group with F149 (Fig. 4g and Extended Data Fig. 7b). The ε-ring at the other end is stabilized mainly by C-H…π interaction of methyl with F171 on ICL3 (see Supplementary Fig. 2 for other possible binding modes of lutein with HeimdallR1). Fucoxanthin fits in the fenestration through the keto end and binds to the protein in a similar fashion as lutein through hydrogen bonding with both WAT331 and T142, as well as π-interaction with F149 (Fig. 4h and Extended Data Fig. 7c). The other possibility, i.e., binding of fucoxanthin in the fenestration via the allene end will not be possible due to steric clash between the acetate moiety on the carotenoid ring and the protein.

Calculated excitation energies reveal a blue-shifted absorption spectrum upon binding of lutein or fucoxanthin to HeimdallR1 (Supplementary Table 3 and Supplementary Fig. 3 for the comparison of excitation energies for different binding modes of lutein). Natural transition orbital analysis (depicting dominant orbitals involved during electronic transitions) reveals that the transitions originate mainly due to local excitations in retinal and lutein/fucoxanthin, and also due to charge transfer from retinal to the respective carotenoid (Extended Data Fig. 8). Electron orbitals are largely localized on the retinal and the extended π-network of the carotenoids backbone with no significant contribution from the rings. This shows that the rings of the carotenoids are important for binding with the rhodopsin but not for energy transfer.

When comparing the calculated structures of HemidallR1 with Kin4B8-XR, the xanthophylls similarly lie transversely against the outer surface of TM6 (Fig. 4i and Extended Data Fig. 7d). However, the angles between the polyene chain and the protein are very different in HeimdallR1 and Kin4B8 due to the helix in ICL3 and the π-bulge in TM6. Moreover, the orientation of the ring is different in Kin4B8 such that its hydroxyl interacts with S208. Most importantly, T142 and F149, which are critical for fucoxanthin and lutein binding in the QM/MM model, are not conserved in Kin4B8 (Fig. 4e). The amino-acid difference in the fenestration may explain the unique binding and energy transfer of fucoxanthin to HeimdallR1.

The rhodopsin family to which HeimdallR1 from Red Sea MAG RS678 belongs has thus far remained an isolated monotypic branch^3,4^. In order to understand the distribution of this family and to assist resolution of its evolutionary ties, we searched multiple databases for similar sequences which yielded three groups of closely related proteins: HeimdallR1-like, HeimdallR2-like and HeimdallR3, found in diverse localities around the globe (Fig. 5a, Extended Data Fig. 9). The HeimdallR1- and HeimdallR2-like clades were closest to each other with pairwise identities ranging between 81% and 86% and all MAGs containing them belonged to the uncultured archaeal species “*Ca.* Kariarchaeum pelagium” from marine and estuarine samples. Interestingly, while nearly all “*Ca.* K. pelagium” genomes had one HeimdallR gene each, phylogenetic relationships between the genomes revealed a branching pattern inconsistent with the HeimdallR1/R2 divide, implying that ongoing recombination might be responsible for the mosaic distribution of the two HeimdallR groups. The third group composed of the sole protein HeimdallR3, which was obtained via co-assembly of metagenomic data from the Groves Creek Marsh (Georgia, USA), showed a modest identity of 63-67% to HeimdallR1/R2 providing thus an important addition for the study of diversity in this family. No complete metagenomic bin could be obtained for the scaffold containing HeimdallR3 but two genes found on the same fragment could be used to place it among “*Ca.* Kariarchaeum” as well: as a potentially new species sister to the sediment-dwelling and HeimdallR-less FT_008 (Fig. 5b). Analysis of the global distribution of HeimdallRs revealed it to be a minor (0-24%) yet widespread family among archaeal proton pumps (ACA, ACB and HeimdallRs) reaching peak relative abundance in the Indian Ocean, the Red Sea and in salt marsh samples along the West Atlantic coast (Fig. 5a). Two proteins representing the additional HeimdallR clades, HeimdallR2 from MAG WBC_A_4_184 obtained from the Apalachicola estuary (Florida, USA)^28^ and HeimdallR3 were chosen for closer investigation and expressed to compare their xanthophyll binding properties to HeimdallR1. Sequence alignment and structure prediction revealed that both proteins have fenestrations and HeimdallR2 in particular is structurally very similar to HeimdallR1, yet interestingly HeimdallR3 appears to lack the helix in ICL3 (Extended Data Fig. 6i). Lutein, diatoxanthin, and fucoxanthin were indeed found to bind to both HeimdallR2 and HeimdallR3 (Extended Data Fig. 10). Fluorescence measurements revealed energy transfer from all three xanthophylls to the retinal moiety in HeimdallR2, while for HeimdallR3 a clear fluorescence signal could be observed only with lutein (Extended Data Fig. 10). We conclude that all pelagic “*Ca.* Kariarchaeum” possess rhodopsin proton pumps capable of utilizing xanthophyll antennas. Gene content analysis of the available genomes, revealed nevertheless a very limited set of genes for carotenoid biosynthesis putatively only capable of synthesis of linear carotenoids of the bacterioruberin series (Extended Data Fig. 10). This indicates that both the retinal and the xanthophyll antennas are obtained by these archaea from the environment. Curiously, other members of the “*Ca.* Kariarchaeaceae” lacked proton-pumping rhodopsins and were restricted to deep-sea and subsurface habitats. Phylogenetic relationships among the “*Ca.* Kariarchaeaceae” and their ties to other heimdallarchaeia point to a singular origin of the pelagic and putatively photoheterotrophic lifestyle of “*Ca.* Kariarchaeum” derived from an ancestral dark habitat (Fig. 5b).

**Fig. 5.**
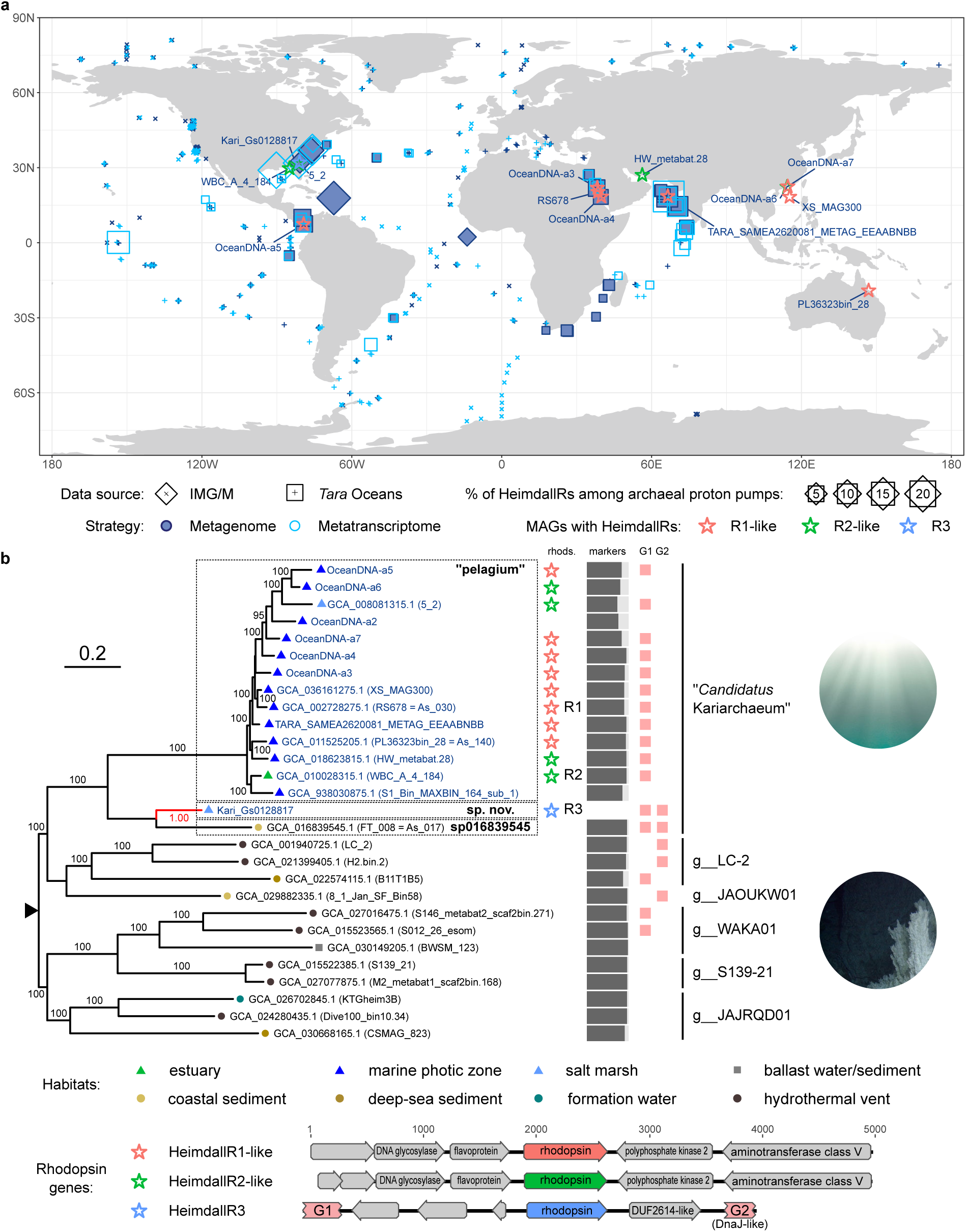
Distribution of diverse marine “*Ca.* Kariarchaeaceae” with HeimdallRs and the phylogenetic relationships between members of “*Ca.* Kariarchaeaceae”. **a,** Global distribution of heimdallarchaeial rhodopsins. Samples in which HeimdallR were detected are indicated with squares with their size proportional to their percentage among all marine archaeal proton pumps. Samples considered but lacking HeimdallRs are indicated with crosses. Locations from which MAGs with HeimdallRs and the HeimdallR3 scaffold (Kari_Gs0128817) originate are labeled and indicated with stars. **b**, Top: Phylogenetic relationships between members of the family “*Ca.* Kariarchaeaceae” based on concatenated alignment of 153 markers present in at least 60% of the genomes. The tree is midpoint-rooted with the root position in agreement with the family root in GTDB (indicated with a triangle). Rapid bootstrap support values are indicated for branches with support >90. The scaffold Kari_Gs0128817 was placed on the phylogenetic tree and its position is indicated in red with the corresponding likelihood weight ratio. Genomes with HeimdallR genes are indicated with stars colored by the corresponding rhodopsin subclade (see Extended Data Fig. 9). Histograms reflect the incidence of phylogenetic markers in the final alignment for each genome and pink squares mark presence of the two genes used for phylogenetic placement of Kari_Gs0128817. Bottom: genomic context of the HeimdallR genes from the three clades. Genes used to place Kari_Gs0128817 on the tree are highlighted.

## Concluding Remarks

Two different spectral tuning strategies are currently known in oceanic microbial rhodopsins; the first is determined mainly by changes in a single amino acid at position 105 (leucine^29,30^ and methionine^31^ in green-absorbing PRs (GPRs) and glutamine^29,30^ in blue-absorbing PRs (BPRs)) and enables PRs to absorb light according to depth, as blue light penetrates deeper in clear oceanic waters^23^. The second strategy is the use of carotenoid antennas absorbing blue light for energy transfer to GPRs, XRs^2^ and HeimdallRs (this study) which otherwise absorb in green. Our present findings, illustrating the advantageous nature of rhodopsin/xanthophyll complexes under white-light illumination, suggest that this strategy might more accurately simulate the solar conditions encountered by marine microbes within aquatic ecosystems (*i.e.* multichromatic light availability at most depths in the photic zone^23^.

We discovered that heimdallarchaeia rhodopsins (HeimdallRs) from “*Ca.* Kariarchaeum”, thus far unique among microbial rhodopsins, can utilize such dissimilar xanthophyll antennas as lutein and fucoxanthin. These pelagic Asgard archaea do not appear to possess genes necessary for synthesis of retinal or any xanthophylls and it is thus possible that this lack of specificity might reflect a spectrum of xanthophyll antennas recruited by them from the environment. Understanding of the exact mechanism by which the tripartite opsin-retinal-xanthophyll complex is assembled will be achieved with the eventual isolation of “*Ca.* Kariarchaeum” in culture.

## Methods

### Proteins chosen for expression

Three representatives from the HeimdallR family were selected based on protein sequence dissimilarity: HeimdallR1 (GenBank accession number MBS85746.1) from metagenomically assembled genome (MAG) RS678 (*Archaea*, “*Ca.* Asgardarchaeota”, “*Ca.* Heimdallarchaeia”, order UBA460, “*Ca.* Kariarchaeaceae”, “*Ca.* Kariarchaeum pelagium”) from the Red Sea (*Tara* Oceans^32^), HeimdallR2 (GenBank accession number NDB54273.1) from MAG WBC_A_4_184 (“*Ca.* Kariarchaeum pelagium”) from the estuary of the Apalachicola River, Florida^28^, and HeimdallR3 from “*Ca.* Kariarchaeum” scaffold Gs0128817_NODE_76 obtained from metagenomic data from Groves Creek Marsh, Skidaway Island, Georgia (JGI project Gs0128817). HeimdallR3 was first identified in the metatranscriptomic fragment Ga0182069_1327250 and the complete ORF could be recovered by recruiting raw metagenomic and metatranscriptomic reads from read runs included in the project with bowtie2 v. 2.3.5.1^33^ using Ga0182069_1327250 as seed, and assembling the recruited reads with spades v. 3.15.5. The resulting contig was further extended in both directions by iterative read recruitment with bowtie2. The highest coverage for the final scaffold was obtained from the metagenomic run SRR7152995 and metatranscriptomic run SRR6980959. ACB-G35 (GenBank accession number MBA4694301.1) from MAG MCMED-G35 (*Archaea*, “*Ca.* Thermoplasmatota”, “*Ca.* Poseidoniia*”*, “*Ca.* Poseidoniales”, “*Ca.* Poseidoniaceae”, species MGIIa-K1 sp003602415)^34^ was selected as a representative of the Archaea clade B family.

### Genome phylogeny

Genomes chosen for phylogenetic reconstruction of the “*Ca.* Kariarchaeaceae” were obtained from GenBank, OceanDNA^35^ and Ocean Microbiomics Database^36^ based on the taxonomic assignment in GTDB r. 220^37^ and source databases. To exclude contaminating scaffolds, a pangenome for “*Ca.* K. pelagium” was obtained with SuperPang v. 1.3.0^38^ (using a relaxed identity threshold of 80% and a k-mer size of 101) and scaffolds contributing to the core pangenome from the individual assemblies were selected. Genes were predicted with Prodigal v. 2.6.3^38^ based on the filtered assemblies (for “*Ca.* K. pelagium”) or entire assemblies (other members of the family). The predicted amino-acid sequences were used to reconstruct phylogeny with PhyloPhlAn v. 3.02^39^ with the “phylophlan” marker database, USEARCH v. 11.0.667 for marker identification, MAFFT v. 7.475^40^ for alignment, TrimAl v. 1.4.1^41^ for alignment trimming and RAxML v. 8.2.12 for phylogenetic reconstruction. Only markers appearing in at least 60% of the genomes were included. The scaffold containing HeimdallR3 was placed on the resulting tree with pplacer v. 1.1.alpha19^42^ using two genes found suitable for this task (homologs found in at least four other “*Ca.* Kariarchaeaceae” genomes with BLASTp from NCBI BLAST+ v. 2.15.0^43^ (E-value threshold of 1e-15): gene G1 of unknown function and G2 coding for a DnaJ-like protein.

### Gene phylogeny

Rhodopsin sequences representative of the different families in the PR-XR superclade were aligned with MAFFT (--auto) and trimmed with TrimAl (-gt 0.9). NeighborNet network was obtained with SplitsTree v. 4.17.0^44^ based on uncorrected p distances. A distance-based tree of the HeimdallR family was obtained by collecting HeimdallR protein sequences at least 210 residues in length, clustering them at 100% identity level with CD-HIT v. 4.8.1^45^, aligning with MAFFT (automatic mode) and reconstructing a neighbor-joining phylogeny in MEGA v. 10.2.5^46^ under Dayhoff model with gamma-distributed substitution rates (shape parameter of 1.0) with 100 bootstrap replicates.

### Global distribution of HeimdallRs

Ocean Microbial Reference Catalog v. 2 (OM-RGC v.2)^47^ and proteins assigned to pfam01036 family in aquatic metagenome and metatranscriptomes assemblies in JGI Integrated Microbial Genomes & Microbiomes (IMG/M)^48^ were used to extract quantitative data on the distribution of HeimdallRs and other marine archaeal proton pump clades. Sequences belonging to the three target clades (HeimdallRs, Archaea clade A and Archaea clade B) were identified by searching the databases with BLASTp using representatives sequences from the PR-XR superclade with an E-value threshold of 1e-10, minimum identity of 50% and a bitscore threshold of 90. Proteins with best matches to one of the three families were assigned to taxa with mmseqs2 v. 14.7e284^49^ using a pre-built database of taxonomically classified rhodopsin sequences extracted from GTDB representative genome assemblies. Sequences classified to *Archaea* were retained. Length-adjusted abundances for the genes were taken directly from the corresponding gene atlas in the case of OM-RGC v. 2 and approximated by dividing the scaffold read depths by scaffold lengths in the case of IMG/M. Per-family abundances were calculated as a sum of the abundance of the individual genes in a sample and relative abundances of HeimdallRs were obtained by dividing their abundance to the total abundance of the three archaeal proton pump clades.

### Carotenoids

Lutein (PHR1699), β-carotene (PHR1239) and fucoxanthin (F6932) were purchased from Sigma-Aldrich. Diatoxanthin was obtained from *Phaeodactylum tricornutum* strain CCAP 1055/5 (Culture Collection of Algae and Protozoa—Scottish Association for Marine Science, Scotland, UK). The extract from dried biomass of *P. tricornutum* was obtained with 80% ethanol. For the separation of diatoxanthin from the extract of *P. tricornutum* biomass, a high performance countercurrent chromatography (HPCCC) system was used in a two-step procedure. The mobile phase consisted of the lower phase of a two-phase solvent system (*n*-heptane, ethyl acetate, ethanol and water in a ratio of 5:4:5:3, *v*/*v*/*v*/*v*), while the stationary phase was the upper phase. In the first HPCCC separation step, an amount of 120 mg of the algal extract was processed with the HPCCC using the mobile phase at a flow rate of 3 mL/min, resulting in 0.25 mg of the fraction containing diatoxanthin. The same separation process was repeated 10 times, and the collected target fractions were pooled, finally yielding 2.5 mg of the diatoxanthin fraction after removal of the solvent by rotary evaporation under reduced pressure at 28°C. To increase the purity of the obtained fraction (2.5 mg), a second HPCCC separation step was performed at a mobile phase flow rate of 1.5 mL/min, yielding in 1.5 mg of the diatoxanthin fraction. The diatoxanthin fraction obtained by the two-step HPCCC was finally purified by gel permeation chromatography using Sephadex LH-20 gel and a mobile phase of 100% methanol. The collected diatoxanthin fraction was evaporated using a rotary evaporator under reduced pressure at 28 °C and yielded 0.57 mg of the compound.

### Mediterranean Sea chromophore extract

450 L of water were sampled on 25 January 2023 at 8 am in the Mediterranean Sea near Michmoret harbor (32° 32.410144 N, 34° 34.846542 E). The water sample was then filtered on a 0.22 µm Durapore® PVDF membrane filter (Milliporem, GVWP14250) after pre-filtration through a mesh net. The sample-containing membranes were then freeze-dried using a lyophilizer (Coolsafa 110-4, ScanVac) for approximately 48 h. Chromophore extraction was done directly on the dried membranes using hexane extraction^50^. Briefly, dried samples were resuspended in 10 mL acetone by applying extensive pipetting and vortexing. Hexane and 10% NaCl were added to the mixture in a 2:2:1 ratio (Acetone: Hexane: 10% NaCl). The mixture was vortexed and then centrifuged at 3,000*g* at 4°C for 3 min. The hexane (top) layer was then transferred to a separate falcon and the process was repeated till the hexane phase became colorless. Combined hexane fractions were then dried using N_2_ gas and either reconstituted in 1 mL Abs ethanol or lyophilized for further characterization.

N_2_-dried lyophilized extracts (10 mg) were resuspended in 1 ml of methanol, agitated for 2 min in 2-ml screw-top polystyrene tubes with 0.5 g of 0.5-mm glass beads under N_2_ in a Genie disruptor and incubated overnight at −20 °C. The chromatographic analysis of the pigments in the extracts was performed in a Merck Hitachi HPLC equipped with a diode array detector according to the method described by^51^. The column used was an RP-18, the flow rate was 1 mL min^−1^, and 100 µL of the sample was injected. The mobile phases used were: Solvent A (ethyl acetate 100%) and solvent B (acetonitrile:H_2_O; 9:1 v/v). The gradient applied was: 0–16 min 0–60% A; 16–30 min 60% A; and 30–35 min 100% A. Standards were supplied by MERCK-SIGMA (Burlington, Massachusetts, United States) or DHI (Hoersholm, Denmark).

### Expression of HeimdallR family representatives

The genes of HeimdallR2 and HeimdallR3 were first optimized for *Escherichia coli* expression system and cloned into pET21a (+) vector with a C-terminal six-His tag, using NdeI and XhoI restriction sites. The point mutation in HeimdallR1 G141F, prepared to block the fenestration in HeimdallR1, was obtained using the NEB Q5 site-directed protocol (https://nebasechanger.neb.com/) with primers 5′- TGTGGTTTGGtttACCCTGAGCGGC-3′ and 5′- CGCATGCCGTCAACG-3′.

*E. coli* C43(DE3) cells harboring the pET21a (+) HeimdallR1 / HeimdallR2 / HeimdallR3 / HeimdallR1-G141F cloned plasmid were grown overnight at 37 °C in LB medium supplemented with ampicillin (50 μg/mL). The day after, the overnight culture was inoculated at a 1:20 dilution into M9 medium containing 50 μg/mL ampicillin. This was grown at 220 rpm and 37 °C until OD_600_= ∼0.6. The expression protein was then induced with 0.25 mM isopropyl β-D-thiogalactopyranoside (IPTG) in the presence of 10 μM all-*trans*-retinal (Toronto Research Chemicals, Canada) at 37 °C for 4 h.

### Expression of ACB-G35 rhodopsin

The gene for ACB-G35 rhodopsin was first optimized for *E. coli e*xpression system and cloned into pET21a (+) vector with a C-terminal six-His tag, using NdeI and XhoI restriction sites.

*E. coli* C43 cells harboring the pET21a (+) ACB-G35 rhodopsin cloned plasmid were grown at 37 °C in LB medium supplemented with ampicillin (50 μg/mL) overnight. The day after, the overnight culture was inoculated at a 1:100 dilution in LB medium containing 50 μg/mL ampicillin. This was grown at 220 rpm and 37 °C until OD_600_= ∼0.6. The expression protein was induced by 1mM IPTG in the presence of 10 μM all-*trans*-retinal (Toronto Research Chemicals, Canada) at 37 °C for 4 h.

### Rhodopsins purification

The rhodopsin-expression *E. coli* cultures were centrifuged at 5,000*g* for 15 min at 4 °C and kept in −80 °C for overnight. The pellet was thawed on ice and resuspended in a buffer containing 50 mM Tris HCl pH 8.0, 5 mM MgCl_2_ and 0.1 mM PMSF (P7626, Sigma-Aldrich). The sample was disrupted by using a microfluidizer for 10 passes at 60 psi. Then, the sample was centrifuged at 5,000*g* for 15 min at 4 °C to pellet undisrupted cells or large cell debris. Membranes were collected by centrifuging the sample at 37,000*g* for 1.5 h at 4 °C and resuspended in a buffer containing 50 mM MES-NaOH pH 6.0, 300 mM NaCl, 5 mM imidazole, 5 mM MgCl_2_, 10% glycerol and 2% DDM final concentration. The sample was incubated overnight at 4 °C with gentle rotation and a second centrifugation at 37,000*g* for 1.5 h at 4 °C was performed. The supernatant was then incubated with Ni-Beads (31103, Cube Biotech) for 1 h. Beads were washed on a gravity column using a buffer containing 50 mM MES-NaOH pH 6.0, 300 mM NaCl, 10% glycerol, 0.05% DDM and 5 mM imidazole. Protein was eluted from the column using a buffer containing 50 mM MES-NaOH pH 6.0, 300 mM NaCl, 10% glycerol, 0.05% DDM and 250 mM imidazole. Eluted protein was washed using Amicon 3-kDa cut-off (UFC800324, Millipore) with storage buffer (50 mM MES-NaOH pH 6.0, 300 mM NaCl, 10% glycerol and 0.05% DDM). The protein was then flash-frozen in liquid N_2_ and stored at −80 °C.

### Binding of environmental chromophore extract to rhodopsin proteins

1mg purified rhodopsin was mixed with the Mediterranean Sea chromophore extract in a ratio of at least 1:3 OD to volume and incubated overnight with gentle rotation at 4 °C. Ethanol volume was adjusted to ensure it did not exceed 1% of the rhodopsin solute volume. Subsequently, Approximately 0.5 mL of Ni-Beads were added to the mixture for 1 h at 4 °C. The rhodopsins were then washed extensively using a storage buffer and eluted using the same buffers used for the initial purification.

### Binding of commercial xanthophyll to rhodopsin proteins

The purified rhodopsin was mixed with carotenoids fresh stock dissolved in dimethyl sulfoxide (DMSO) and incubated overnight with gentle rotation at 4 °C. The molar quantities of each component were estimated based on their respective absorption spectra, utilizing molar extinction coefficients of ε = 45,000-50,000 M^-1^ cm^-1^ at the absorption peak of rhodopsin and ε = 145,100 M^-1^ cm^-1^ at 445 nm for carotenoids. The volume of DMSO was adjusted to ensure it did not exceed 5% of the solution volume of rhodopsin. Subsequently, approximately 0.5 mL of Ni-Beads were added to the mixture for 1 h at 4 °C. The rhodopsins were then washed extensively using a storage buffer and eluted using the same buffers used for the initial purification.

### Absorption spectroscopic measurements

Absorption spectral measurements of HeimdallR1, HeimdallR2, HeimdallR3, ACB-G35 and Kin4B8-XR with and without carotenoids were taken with a Shimadzu (UV-1800) Spectrophotometer. The absorption spectral measurements of HeimdallR1 and ACB-G35 with carotenoid extract were taken with a BioTek Synergy MX Plate Reader.

### Fluorescence spectroscopic measurements

Fluorescence emission and excitation spectral measurements were taken on a Jobin Yvon-Spex Fluorolog-3 spectrofluorometer. The spectrofluorometer is equipped with 450W Xe-lamp as the light source, double-grating monochromator in the excitation, single-grating in emission positions, and a photomultiplier tube detector (R928P). All measurements were done at pH 5.5 and in slit width at 10 nm in the excitation and emission. Emission was monitored at 720 nm and excitation between 400 nm to 650 nm. The absorbance of all samples was kept at OD < 0.5. Fluorescence excitation spectral profile was corrected by the absorbance spectra profile for each nm, due to the inner filter effect: *F_ideal_* = *F_flu_*10^(*A_ex_ + A_em_*)^/2, where *F_ideal_* is the ideal fluorescence intensity after we consider the inner filter effect, *F*_*flu*_ is the fluorescence intensity measured, *A_ex_* is the absorbance at the *F*_*flu*_ excitation point and *A*_*em*_ is the absorbance at the emission point (always at 720 nm in our case)^52^.

### Protein expression and purification of HeimdallR1 for photocycle measurements

*E. coli* C43(DE3) cells harboring the HeimdallR1-cloned plasmid were grown at 37 °C in LB medium supplemented with ampicillin (50 μg/mL) overnight. The day after, the overnight culture was inoculated in M9 medium containing 50 μg/mL ampicillin. This was grown at 200 rpm and 37 °C (until OD_600_= ∼0.6). The expression of C-terminal 6x His-tagged proteins was induced by 0.1 mM IPTG in the presence of 10 μM all-*trans*-retinal (Toronto Research Chemicals, Canada) at 37 °C for 4 h. The harvested cells were sonicated (Ultrasonic Homogenizer VP-300N; TAITEC, Japan) for disruption in a buffer with 50 mM Tris–HCl (pH 8.0) and 5 mM MgCl_2_. The membrane fraction was collected through ultracentrifugation (CP80NX; Eppendorf Himac Technologies, Japan) at 142,000*g* for 1 h. The proteins were solubilized in a buffer containing 50 mM MES–NaOH (pH 6.5), 300 mM NaCl, 5 mM imidazole, 5 mM MgCl_2_, and 3% *n*-dodecyl-β-D-maltopyranoside (DDM) (ULTROL Grade; Calbiochem, Sigma-Aldrich, MO). Solubilized proteins were separated from the insoluble fractions through ultracentrifugation at 142,000*g* for 1 h. Proteins were purified using a Co-NTA affinity column (HiTrap TALON crude; Cytiva, MA). The resin was washed with a buffer containing 50 mM MES–NaOH (pH 6.5), 300 mM NaCl, 50 mM imidazole, 5 mM MgCl_2_, and 0.1% DDM. Proteins were eluteined in a buffer containing 50 mM Tris–HCl (pH 7.0), 300 mM NaCl, 300 mM imidazole, 5 mM MgCl_2_, and 0.1% DDM. Eluteined protein was immediately concentrated using a 50 mL centrifugal ultrafiltration filter with a molecular weight cutoff of 30-kDa (Amicon Ultra-4, Millipore) and the buffer exchanged with buffer 50 mM HEPES–NaOH pH 7.0, 150 mM NaCl, 10% glycerol, and 0.1% DDM.

### Purification of carotenoid-binding HeimdallR1

Carotenoid-binding HeimdallR1 was obtained by mixing purified HeimdallR1 with carotenoids (lutein or fucoxanthin). First, carotenoids were dissolved in dimethyl sulfoxide (DMSO) at twice the molar amount of HeimdallR1 to be bound. The molar amounts of each were estimated by measuring the absorption spectrum (Extended Data Fig. 3a), using a molar extinction coefficient of *ε* = 45,000 M^−1^ cm^−1^ at the absorption maximum of rhodopsin and *ε* = 145,100 M^−1^ cm^−1^ at 445 nm for carotenoids. The volume of DMSO was adjusted so that it did not exceed 2% of the rhodopsin solution volume. After preparing the carotenoid solution as described above, it was added to HeimdallR1 and left on ice and in the dark for 2 h, gently mixing the solution every 15 min. To remove the excess of carotenoids not bound to HeimdallR1, it was purified again using the same method as the purification of the original HeimdallR1 protein. The purified protein was immediately concentrated using a 50 mL centrifugal ultrafiltration filter with a molecular weight cutoff of 30-kDa (Amicon Ultra-4, Millipore) and the buffer exchanged with buffer 20 mM HEPES–NaCl (pH 7.0), 100 mM NaCl, 0.05% DDM.

### Laser-flash photolysis

For the laser-flash photolysis spectroscopy, HeimdallR1 with/without lutein or fucoxanthin was solubilized in 20 mM HEPES–NaCl (pH 7.0), 100 mM NaCl, 0.05% DDM. Optical density (OD) of the rhodopsin was adjusted to approximately 0.4–0.5 (protein concentration of approximately 0.2–0.25 mg mL^−1^) for HeimdallR1 and HeimdallR1 with fucoxanthin and approximately 0.2 (protein concentration of approximately 0.1 mg mL^−1^) for HeimdallR1 with lutein at the λ^a^_max_. The laser-flash photolysis measurement was conducted as previously described^4,53^. Nano-second pulses from an optical parametric oscillator (λ_exc_ = 550 (without lutein), 535 (with lutein), and 540 nm (with fucoxanthin), 4.5 mJ pulse^−1^ cm^−2^, 1.1 Hz (basiScan, Spectra-Physics, CA) pumped by the third harmonics of Nd-YAG laser (λ = 355 nm, INDI40, Spectra-Physics, CA) were used for the excitation of HeimdallR1 w/wo lutein and w/wo fucoxanthin). The transient absorption spectra were obtained by monitoring the intensity change of white-light from a Xe-arc lamp (L9289-01, Hamamatsu Photonics) passed through the sample with an ICCD linear array detector (C8808-01, Hamamatsu Photonics). To increase the signal-to-noise ratio, 40–60 spectra were averaged, and the singular-value decomposition analysis was applied. To measure the time evoluteinion of transient absorption change at specific wavelengths, the output of a Xe-arc lamp (L9289-01, Hamamatsu Photonics) was monochromated by monochromators (S-10, Soma Optics) and the change in the intensity after the photoexcitation was monitored with a photomultiplier tube (R10699, Hamamatsu Photonics). To increase signal-to-noise ratio, 200–400 signals were averaged. The signals were globally fitted using a multi-exponential function to determine the lifetimes and absorption spectra of each photointermediate.

To measure the fold change in transient absorption change between with and without xanthophylls, nanosecond pulses from an optical parametric oscillator (basiScan, Spectra-Physics) pumped by the third harmonics of Nd–YAG laser (λ = 355 nm, INDI40, Spectra-Physics, CA) were used for the excitation of HeimdallR1 at different wavelengths (λ_exc_ = 425, 450, 465, 480, 545, and 590 nm for lutein, and λ_exc_ = 440, 465, 580, 490, 530, 540, 550, and 600 nm for fucoxanthin). The pulse energy was adjusted to 0.55 mJ cm^−2^ and 0.56 mJ cm^−2^ for lutein and fucoxanthin, respectively, to keep the linearity between the number of the absorbed photons and the transient absorption change. QY based on fold change in transient absorption change between with and without xanthophylls were calculated using the following equations:

For lutein:

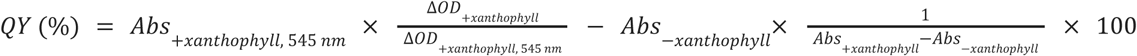

For fucoxanthin:

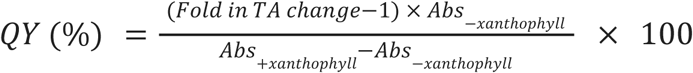

### High-performance liquid chromatography (HPLC) analysis of retinal isomers

Retinal configuration was analyzed by HPLC using purified HeimdallR1, HeimdallR1 with lutein or fucoxanthin in a buffer containing 20 mM HEPES–NaCl (pH 7.0), 100 mM NaCl, 0.05% DDM. Before the measurements, the OD of the rhodopsin was adjusted to approximately 0.2 (protein concentration of approximately 0.1 mg mL^−1^), and the proteins were stored at 4 °C overnight in the dark for dark-adapetd (DA) samples. The HPLC system was equipped with a silica column particle size of 3 μm, 150 × 6.0 mm (Pack SIL, YMC, Japan), pump (PU-4580, JASCO, Japan), and UV–vis detector (UV-4570, JASCO, Japan). The solvent was composed of 15% (v/v) ethyl acetate and 0.15% (v/v) ethanol in hexane and with a flow rate of 1.0 mL min^−1^. To denature the protein, 280 μL of 90% methanol solution was added to the 75 μL sample. Retinal oxime formed by the hydrolysis reaction with 25 μL of 2 M hydroxylamine solution was extracted with 800 μL of hexane, and 200 μL of the solution was injected into the HPLC system. For measurements during light illumination (Light), the sample solutions were illuminated at λ = 550 ± 10 nm for HeimdallR1 and HeimdallR1 with fucoxanthin and at λ = 540 ± 10 nm fro HeimdallR1 with lutein (Bandpass, AGC Techno Glass, Japan) for 1 min, followed by denaturation and hydrolysis of the retinal chromophore under illumination. For measurements of light-adapted (LA) samples, the sample solution was illuminated at λ = 550 ± 10 nm for HeimdallR1 and HeimdallR1 with fucoxanthin and at λ = 540 ± 10 nm for HeimdallR1 with lutein for 1 min, and after waiting for 1 min, denaturation and hydrolysis reactions of the retinal chromophore were conducted. The molar compositions of the retinal isomers were calculated from the areas of the corresponding peaks in the HPLC patterns. The molar composition of the retinal isomers in the sample was determined with the molar extinction coefficients at 360 nm for each isomer (all-*trans*-15-*syn*: 54,900 M^−1^ cm^−1^; all-*trans*-15-*anti*: 51,600 M^−1^ cm^−1^; 13-*cis*-15-*syn*, 49,000 M^−1^ cm^−1^; 13-*cis*-15-*anti*: 52,100M^−1^ cm^−1^; 11-*cis*-15-*syn*: 35,000 M^−1^ cm^−1^; 11-*cis*-15-*anti*: 29,600 M^−1^ cm^−1^). Three independent measurements were performed to estimate experimental error.

### pH titration

To investigate the pH dependence of the absorption spectra of HeimdallR1 with/without lutein and with/without fucoxanthin, the OD of the rhodopsin was adjusted to approximately 0.5 (protein concentration of approximately 0.25 mg mL^−1^) and solubilized in a 6-mix buffer (trisodium citrate, MES, HEPES, MOPS, CHES, CAPS (10 mM each, pH 7.0), 100 mM NaCl, and 0.05% DDM). The pH was adjusted to the desired value by the addition of small aliquots of 1-5 N HCl and NaOH. Absorption spectra were recorded using a UV–vis spectrometer (V-750, JASCO, Japan). The measurements were performed at every 0.3–0.6 pH values.

### Low-temperature UV-visible and FTIR spectroscopic analysis

Samples for low-temperature UV-visible and FTIR spectroscopy were prepared as described for photocycle measurements with small modifications. The proteins were solubilized by the use of 1 % DDM (Anatrace, USA). Then, the samples, HeimadallR1 with lutein, with fucoxanthin, or without xanthophylls, were reconstituted into POPE:POPG membrane as described previously^2^.

Low-temperature UV-visible spectroscopy and FTIR spectroscopy were performed as described previously^2^. Briefly, lipid reconstituted HemidallR1 was placed onto a BaF_2_ window making dry films. Hydrated films with 1 µL H_2_O or D_2_O were fixed to cryostat (Optistat, Oxford Instruments) attached to UV-visible (V-750, JASCO) and FTIR (Cary670, Agilent) spectrometers. K intermediate of HemidallR1 was generated by irradiating 530 nm light for 30 s using an interference filter (KL53, Toshiba) at 77 K, which reverted to the original state by >590 nm for 30 s with a cut-off filter (R-61 cut-off filter, Toshiba). Light activated difference spectra were obtained by subtracting spectra before light irradiation from after light irradiation at 77 K. Averages of 100 experiments were conducted for the spectra of HeimadallR1 with lutein, with fucoxanthin, or without xanthophylls.

### Proton-pumping measurements

The protocol was modified from^54^. In brief, 200 ml of rhodopsin-expressing *E. coli* cells was centrifuged at 3,600*g* for 10 min and resuspended into 20 ml of 30 mM Tris-HCl pH 8.0 and 20% sucrose. 200 µg of lysozyme (L6876, Sigma-Aldrich) was added to the cell suspension and gently rotated for 1 h at room temperature. Spheroplasts were centrifuged at 3,600*g* for 15 min at room temperature and the pellet was resuspended with 6 ml of unbuffered solution (10 mM NaCl, 10 mM MgSO_4_·7H_2_O and 100 μM CaCl_2_). Addition of 100 µM lutein, diatoxanthin or fucoxanthin to approximately 2 ml (a third of the volume) followed by overnight incubation at 4 °C with gentle rotation. Spheroplasts were then washed three times with unbuffered solution (10 mM NaCl, 10 mM MgSO_4_·7H_2_O and 100 μM CaCl_2_).

Samples were kept in the dark until the pH was stabilized and illuminated using Max-303 compact xenon lamp (Asahi Spectra, Japan). The light intensity at the sample location was 860 µmol m^−2^ s^−1^ for the different carotenoids experiment (Fig. 3a), and 2,000, 860, 590, and 515 µmol m^−2^ s^-1^ for the different light intensity experiment (Fig. 3b). The light intensity measurements were performed using a LI-COR Biosciences LI-250A light meter. pH was monitored using a LAQUA F-72G pH/ION meter (HORIBA scientific, Japan) equipped with a 9618S-10D pH microelectrode. To explore the effect of xanthophyll antennas on proton pump activity, pH results were converted to proton concentration. An initial analysis of the data determined that proton concentrations were changing in a near-linear manner in the range between 40 and 70 seconds after illumination onset and thus the rate of proton pumping was approximated by the slope of the regression line for Δ[H^+^] over time in this range: *ŷ*(*t*) = *a* + *b*· *t*. The experiment was conducted in a pairwise manner: A single preparation of spheroplasts derived from a single *E. coli* colony (all prepared from the same transformation stock expressing the rhodopsin gene) was used to obtain proton pumping rates for the protein with (*b*^*r*+*a*^) and without the carotenoid antenna (*b*^*r*^). In total, at least four independent spheroplast preparations were utilized for each protein assay with and without a carotenoid antenna. Spheroplast samples exhibiting low rhodopsin expression where proton pumping activity was near or below the detection threshold of the pH meter used, were excluded from the analysis. A total of nine samples were excluded due to insufficient proton pumping activity. Fold change was calculated as the ratio between the two rates: *b*^*r*+*a*^ ÷ *b*^*r*^. The resulting log2-fold changes (LFC) were used to analyze the influence of the presence/absence of the fenestration, light intensity and type of antenna using linear models in R v. 4.4.1^55^. Since the same batches of spheroplasts were used to obtain LFC values for different light intensities, the model for the light intensity experiment (with lutein as the antenna) was fit with batch as a random effect in lme4 v. 1.1-31^56^ with the following design formula: *LFC* ∼ *intensity* ∗ *fenestration* + (1|*batch*). For the experiment with varying antenna types (at 860 µmol m^−2^ s^−1^), a simpler linear model was fit: *LFC* ∼ *antenna type* ∗ *fenestration*. Type-II analysis of deviance tables were obtained with the Anova function from the car package v. 3.1-1^57^. Distribution of the residuals was checked visually using Q–Q plots and while they were normally distributed in the light-intensity analysis, outliers in the antenna-type analysis were found to cause deviation from the expected distribution of the residuals. Removal of these outliers substantially improved the fit but did not lead to qualitative changes in the results of the analysis.

### Protein expression and purification for structural analysis

pET21a-HeimdallR1 was transfected in *E. coli* C41 (Rosetta). The transformant was grown in LB supplemented with 50 µg ml^−1^ ampicillin at 220 r.p.m. at 37 °C. When the OD_600_ reached 0.6, expression was induced using a 1 mM IPTG. The induced culture was grown at 120 r.p.m. for 4 h at 37°C, in the presence of 10 µM all-*trans* retinal. The collected cells were disrupted by sonication in buffer containing 20 mM Tris-HCl pH 8.0, 200 mM NaCl and 10% glycerol. The crude membrane fraction was collected by ultracentrifugation at 180,000*g* for 1 h. The membrane fraction was solubilized in buffer containing 20 mM Tris-HCl pH 8.0, 200 mM NaCl, 1.5% DDM, 10% glycerol and 10 mM imidazole for 1 h at 4 °C. The supernatant was separated from the insoluble material by ultracentrifugation at 140,000*g* for 30 min and incubated with Ni-NTA resin (Qiagen) for 30 min. The resin was washed with 10 column volumes of wash buffer containing 20 mM Tris-HCl pH 8.0, 500 mM NaCl, 0.03% DDM, 10% glycerol and 20 mM imidazole. The resin was incubated overnight (more than 12 h) with 2 column volume of wash buffer containing 150 µM fucoxanthin. Then, the resin was washed with 5 column volumes of wash buffer. The protein was eluted in buffer containing 20 mM Tris-HCl pH 8.0, 500 mM NaCl, 0.03% DDM, 10% glycerol and 300 mM imidazole. The eluate was dialysed against buffer containing 20 mM Tris-HCl pH 8.0, 150 mM NaCl and 0.03% DDM. The protein bound to fucoxanthin was concentrated to 50 mg ml^−1^ using a centrifugal filter device (Amicon 10 kDa MW cut-off), and frozen until crystallization.

### X-ray crystallographic analysis of HeimdallR1

Since cryo-electron microscopy did not provide structural data, X-ray crystallographic analysis was performed. The protein bound to fucoxanthin was reconstituted into monoolein at a weight ratio of 1:1.5 (protein to lipid). The protein-laden mesophase was dispensed into 96-well glass plates in 30-nL drops and overlaid with 800 nL precipitant solution, using a Gryphon robot (ARI), as previously described^58^. Crystals of HeimdallR1 were grown at 20 °C in precipitant conditions containing 30% PEG300, 100 mM sodium acetate pH 3.9 and 100 mM sodium nitrate. The crystals were harvested directly from the lipid cubic phase using micromeshes (MiTeGen) and frozen in liquid nitrogen, without adding any extra cryoprotectant.

X-ray diffraction data were collected at the SPring-8 beamline BL32XU with an EIGER X 9M detector (Dectris), using a wavelength of 1.0 Å. Small-wedge (10° per crystal) datasets were collected using a 15 × 10-μm^2^ beam with the ZOO system^59^, an automatic data-collection system developed at SPring-8. The collected images were processed using KAMO^60^ with XDS^61^, and 132 datasets were indexed with the consistent unit cell parameters. After correlation coefficient-based clustering using normalized structure factors followed by merging using XSCALE^62^ with outlier rejections implemented in KAMO, 115 datasets were selected for the downstream analyses, because it gave the highest inner-shell and outer-shell CC_1/2_. The HeimdallR1 structure was determined by molecular replacement with PHASER^62^, using the model calculated by AlphaFold2^20^. Subsequently, the model was rebuilt and refined using Coot^63^ and phenix.refine^64^. The final model of HeimdallR1 contained all the residues (1–246), retinal, 3 monoolein molecules, 2 nitrate ions and 121 water molecules. The fucoxanthin molecule was not modelled, due to the absence of the corresponding density.

### QM/MM simulations of HeimdallR1 with lutein and fucoxanthin

The structures of the carotenoids were docked by visual inspection to HeimdallR1 and by comparison to zeaxanthin-bound Kin4B8 (PDB: 8I2Z)^2^. The structures were further optimized using the hybrid quantum mechanics/molecular mechanics (QM/MM) method^27^. Retinal linked to K231 was considered in the QM region along with lutein or fucoxanthin. A region of 5 Å around the retinal protonated Schiff base and the carotenoid was considered in the active region during optimization. The QM/MM boundary was placed between Cδ and Cε of the lysine sidechain. The QM part was described using Grimme’s GFN2-xTB method^65^ whereas the remaining part of the protein within the active region was treated using the AMBER ff14SB force field^66^ (MM part). The water molecules were described using the TIP3P model^67^. To model non-bonded interactions, the force switching scheme was implemented on the coulomb interaction with a cutoff of 12 Å^68^. Optimization was performed using the L-BFGS method using the ORCA 5.0 program^69^. Excitation energies and natural transition orbitals were computed at the RI-ADC(2)^70^ level in combination with the cc-pVDZ^71^ basis set using Turbomole program package^72^.

## Data availability

All data are available in the main text or the Supplementary Information. Results of the bioinformatic analyses are deposited in the Figshare repository https://doi.org/10.6084/m9.figshare.26906593. Atomic coordinates of the crystal structure of HeimdallR1 have been deposited in the Protein Data Bank under XXX.

## Code availability

The code used for the bioinformatic analyses is available from the GitHub repository (https://github.com/BejaLab/Kariarchaeaceae).

## Acknowledgements.

We thank S. Balashov, M. Sheves and G. Yahel for discussing rhodopsins and oceanography with us, and R. Edrei for help with spectroscopic measurements. O.B. is grateful to the Atmosphere and Ocean Research Institute (U.Tokyo, Japan) for the stay during the work on the manuscript. This work was supported by the European Commission, under Horizon Europe’s research and innovation programme (Bluetools project, Grant Agreement No 101081957 to O.B.), the Nancy and Stephen Grand Technion Energy Program (GTEP, to O.B.), the Israel Science Foundation (Research Center grant 3131/20 to I.S. & O.B.), The German-Israeli Foundation for Scientific Research and Development (GIF NEXUS grant I-1560-207.9/2023 to I.S. & O.B.), Deutsche Forschungsgemeinschaft (DFG SFB 1078 to I.S.), Swiss National Science Foundation (SNF Sinergia grant 213507 to I.S.), the Agencia Estatal de Investigación (grant PID2022-140995OB-C21 by MICIU/AEI/ 10.13039/501100011033 and ERDF/EU to R.L.), the Czech Ministry of Education (OP JAK project Photomachines reg. no. CZ.02.01.01/00/22_008/0004624 to M.Koblížek), the Institute for Fermentation Osaka (W.S.), JSPS KAKENHI Grants-in-Aid (grants JP21H04969 to H.K., JP23H04404 and 24H02268 to K.I., JP19H05777 to W.S., JP23KJ0721 to Y.Matsuzaki, JP24K23232 to T.T., and JP24KJ0909 to S.M.), JST CREST (grants JPMJCR1753 to H.K., JPMJCR22N2 to K.I., and JPMJCR20E2 to O.N.), and MEXT Promotion of Development of a Joint Usage / Research System Project: Coalition of Universities for Research Excellence Program (CURE) (grant JPMXP1323015482 to H.K. and K.I.), the Platform Project for Supporting Drug Discovery and Life Science Research (Basis for Supporting Innovative Drug Discovery and Life Science Research (BINDS)) from AMED, grant numbers JP24ama121002 (support number 3272, O.N.) and JP24ama121012 (supporting number 6234 to K.I.). M.d.C.M. is grateful to the Azrieli Foundation for the award of the Azrieli International Postdoctoral Fellowship (cohort 2022-2023). O.B. holds the Louis and Lyra Richmond Chair in Life Sciences.

## Author contributions

G.T. and A.C. conceived the project. G.T. performed environmental carotenoid extractions and binding to rhodopsins, and measured light-dependent proton pumping activity. M.Konno, R.A.-Y. and M.d.C.M. performed expression and purification of HeimdallR1 and carotenoid-binding HeimdallR1. M.d.C.M. performed HPLC analysis of retinal isomers, pH-titration, and experimental data analysis. M.d.C.M. and K.I. performed laser-flash photolysis. A.R. performed bioinformatics. S.L. and M.Konno performed molecular biology. A.M.-M., and R.L. performed carotenoid characterization from environmental samples and rhodopsin-bound carotenoids. M.Koblížek, D.B.-P. and J.C. purified diatoxanthin from *P. tricornutum*. S.I., Y.Mizonu, K.K. and H.K. performed low-temperature UV–Vis and FTIR spectroscopy. Y.Matsuzaki, T.T., S.M., W.S. and O.N. performed structural analyses. P.N. and I.S. performed MD simulations, hybrid QM/MM simulations and interpreted the results of the simulations. O.B. coordinated the project, and wrote the paper with input from all authors.

## Extended data figures and tables

**Extended Data Fig. 1.**
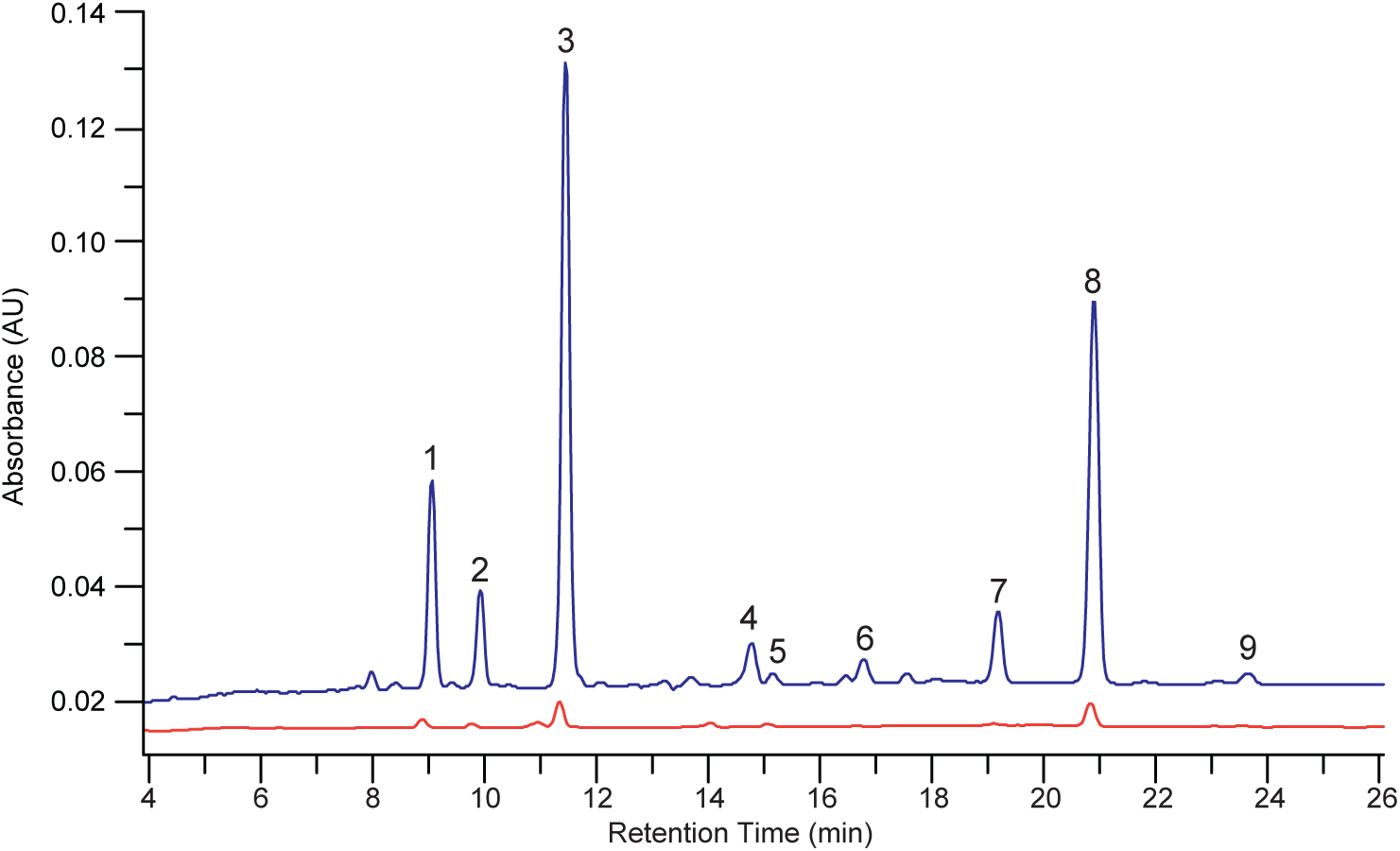
Environmental marine xanthophylls bind to HeimdallR1. HPLC profile of Mediterranean Sea chromophore extract (blue) and HeimdallR1 bound chromophores (red). Main peaks correspond to fucoxanthin (2), diatoxanthin (3), lutein (5), β-carotene (8), and uncharacterized (1,4,6,7,9). Chromophores were registered at 450 nm.

**Extended Data Fig. 2.**
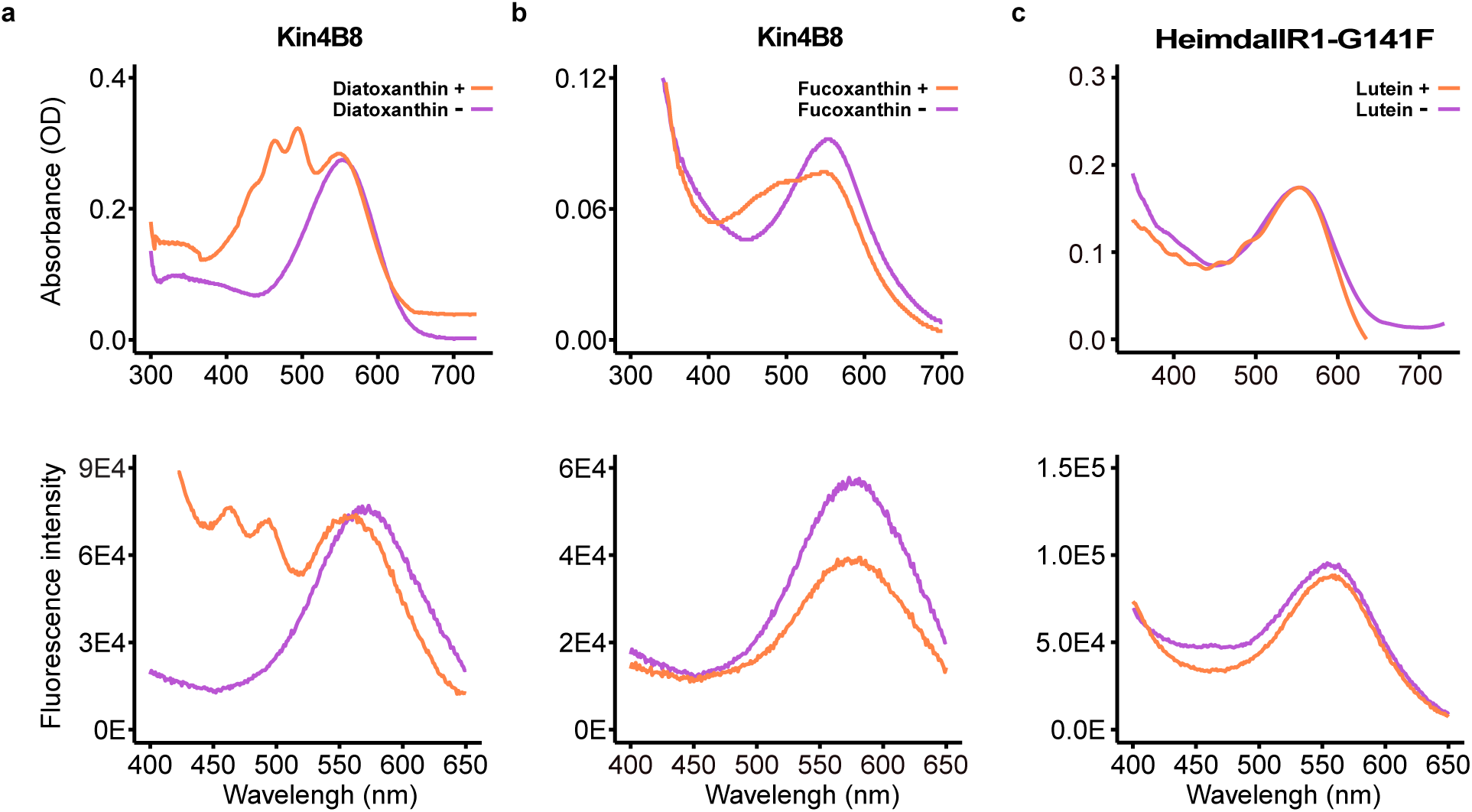
Spectroscopic characterization of Kin4B8-XR and HeimdallR1-G141F with hydroxylated carotenoids. **a**, Absorbance change (up) and fluorescence excitation spectra (bottom) of Kin4B8-XR upon incubation with (orange) or without (purple) diatoxanthin. **b**, Absorbance change (up) and fluorescence excitation spectra (bottom) of Kin4B8-XR upon incubation with (orange) or without (purple) fucoxanthin. **c**, Absorbance change (up) and fluorescence excitation spectra (bottom) of HeimdallR1-G141F upon incubation with (orange) or without (purple) lutein; emissions were recorded at 720 nm.

**Extended Data Fig. 3.**
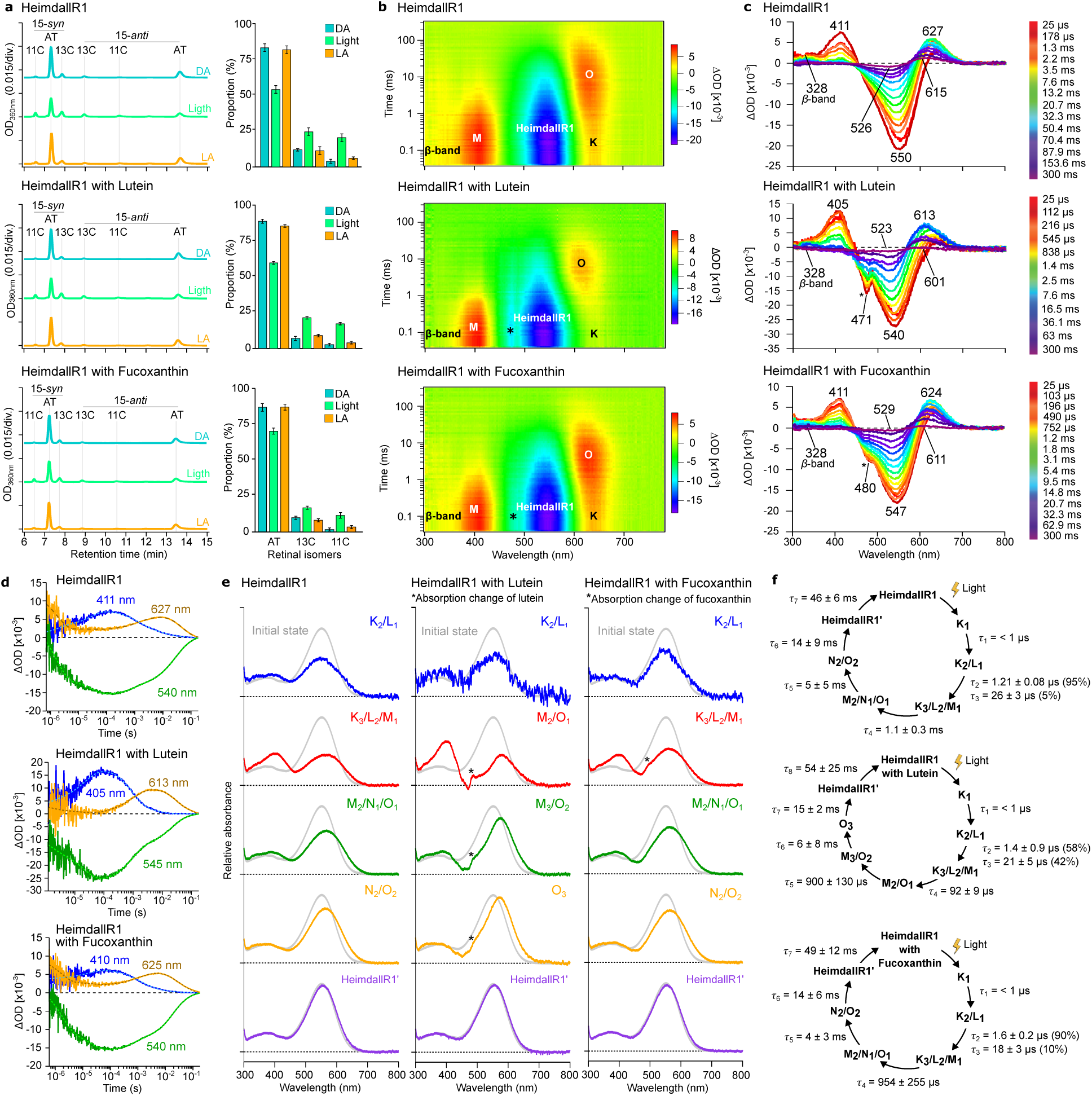
The photocycle of HeimdallR1. **a**, Chromatogram of HPLC analyses (left) and the compositions of the retinal isomers (right) under the dark (DA, blue), light (green), and light-adapted (LA, orange) conditions, where AT, 13C, 11C, syn, and anti indicate all-*trans*, 13-*cis*, 11-*cis*, *syn*, and *anti* configurations, respectively, **b**, two-dimensional plot of transient absorption change, and **c**, transient absorption spectra at different time points of HeimdallR1 without (top), with lutein (center), and with fucoxanthin (bottom). **d**, Time course of the transient absorption change of HeimdallR1 without (top), with lutein (center), and with fucoxanthin (bottom). **e**, Absorption spectra of photointermediates of HeimdallR1 without (left), with lutein (center), and with fucoxanthin (right). **f**, Photocycle models of HeimdallR1 without (top), with lutein (center), and with fucoxanthin (bottom).

**Extended Data Fig. 4.**
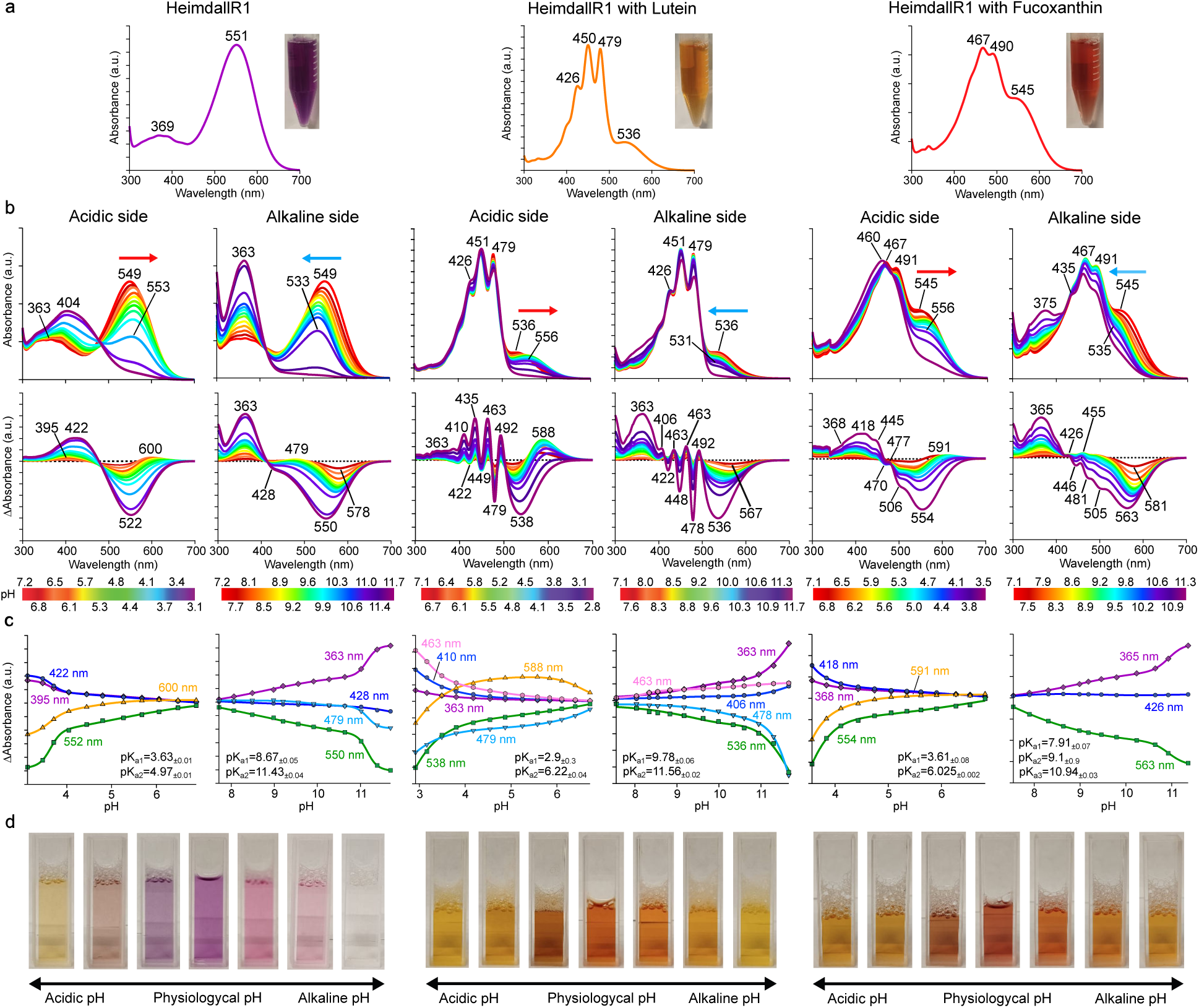
Absorption spectra of purified HeimdallR1 with and without lutein or fucoxanthin and pH dependence on the absorption. **a**, UV–vis absorption spectra. Pictures of purified proteins are shown next to the corresponding results. **b**, Absorption spectra (top) and the different absorption spectra (bottom), and **c**, pH titration curves for the calculation of p*K*_a_ were measured depending on acidic (left) and alkaline (right) pH changes. The red-shifted (red arrow) and blue-shifted (blue arrow) absorption spectra at each pH are indicated by arrows. The pH titration curves were analyzed using the Henderson–Hasselbalch equation^73^. **d**, Color protein changes upon acid–basic titration.

**Extended Data Fig. 5.**
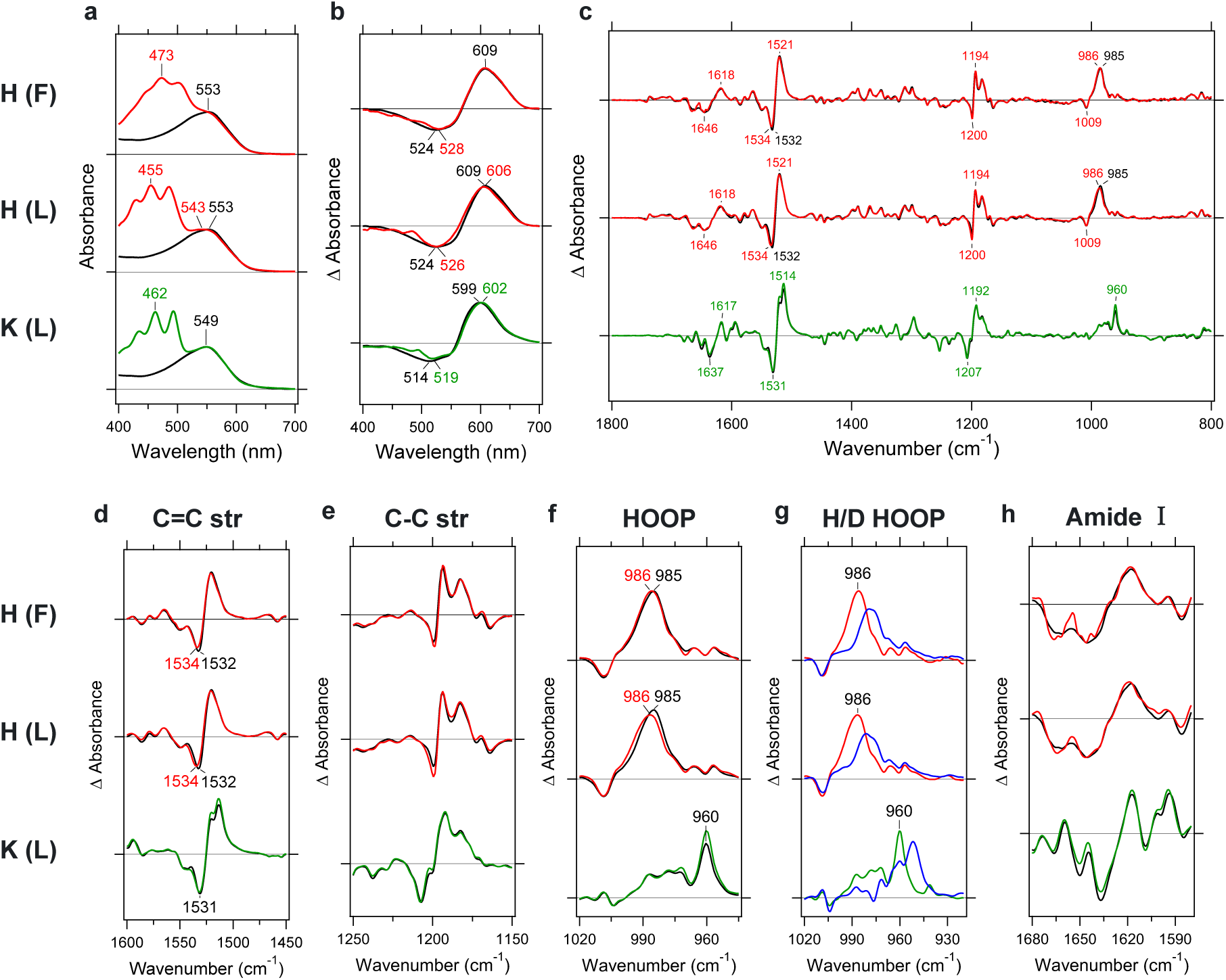
Influence of lutein or fucoxanthin on the retinal photoisomerization in HeimdallR1 at 77 K. UV-visible (**a**, **b**) and FTIR (**c**-**h**) spectra obtained for lipid-reconstituted HeimdallR1 with (red; H(F)) or without (black; H(-)) fucoxanthin (top) and HeimdallR1 with (red; H(L)) or without (black; H(-)) lutein (middle) are compared to those for lipid-reconstituted Kin4B8-XR with (green; K(L)) or without (black; K(-)) lutein (bottom) previously published^2^. **a**, UV-visible absorption spectra of H(F) (red) or H(-) (black) (top), H(L) (red) or H(-) (middle), and K(L) (green) or K(-) (black) (bottom) at 77 K. One division of the y-axis corresponds to 1.0 absorbance units. **b**, Difference UV-visible spectra upon illumination of H(F) (red) or H(-) (black) (top), H(L) (red) or H(-) (black) (middle), and K(L) (green) or K(-) (black) (bottom) at 77 K. Hydrated films of lipid-reconstituted protein were illuminated at 540 nm light, which forms the red-shifted K intermediate. One division of the y-axis corresponds to 0.08 absorbance units. **c**, Light-minus-dark difference FTIR spectra upon illumination of H(F) (red) or H(-) (black) (top), H(L) (red) or H(-) (black) (middle), and K(L) (green) or K(-) (black) (bottom) at 77 K. Hydrated films of lipid-reconstituted protein with H_2_O were first illuminated at 540 nm light (solid lines), which forms the K intermediate, and the K intermediate was then reverted by illumination at >590 nm light (dotted lines). Spectral acquisition was repeated to improve signal-to-noise ratio. Positive and negative signals originate from the K intermediate and unphotolyzed state, respectively. One division of the y-axis corresponds to 0.0032 absorbance units. **d**, Enlarged spectra of the C=C stretching frequency region of the retinal chromophore (1600-1450 cm^-1^) from (**c**). Negative peaks are different between H(F) (1534 cm^-1^) and H(-) (1532 cm^-1^) (top) or between H(L) (1534 cm^-1^) and H(-) (1532 cm^-1^) (middle), but not for K(L) and K(-). One division of the y-axis corresponds to 0.003 absorbance units. **e**, Enlarged spectra of the C-C stretching frequency region of the retinal chromophore (1250-1150 cm^-1^) from (**c**). Spectra are identical with and without xanthophylls. One division of the y-axis corresponds to 0.002 absorbance units. **f**, Enlarged spectra of the hydrogen out-of-plane (HOOP) vibrational region of the retinal chromophore (1020-945 cm^-1^) from (**c**). Positive peaks are different between H(F) (986 cm^-1^) and H(-) (985 cm^-1^) (top) or between H(L) (986 cm^-1^) and H(-) (985 cm^-1^) (middle), but not for the 960-cm^-1^ band (bottom). One division of the y-axis corresponds to 0.0015 absorbance units. **g**, Spectral comparison of the HOOP bands between H_2_O (red or green) and D_2_O (blue) hydrations. Down-shifts of the positive peaks at 986 cm^-1^ (top, middle) and 960 cm^-1^ (bottom) show that these HOOP bands originate from the Schiff base region. One division of the y-axis corresponds to 0.0015 absorbance units. **h**, Enlarged spectra of the amide-I region of the peptide backbone (1680-1575 cm^-1^) from (**c**). Spectra are identical with and without xanthophylls. One division of the y-axis corresponds to 0.001 absorbance units.

**Extended Data Fig. 6.**
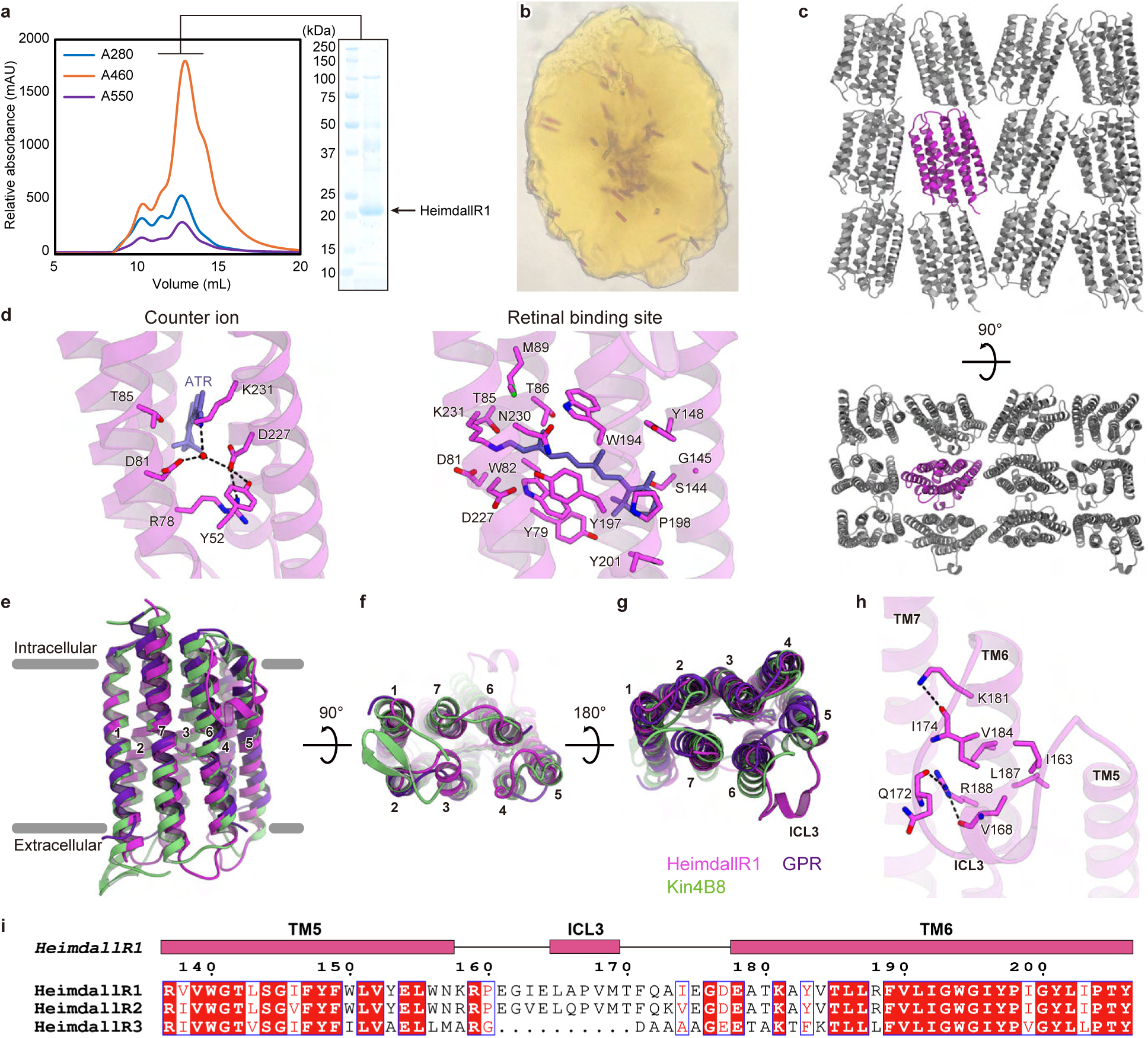
Structural features of HeimdallR1. **a**, Size-exclusion chromatography profile of the HeimdallR1 bound to fucoxanthin. The relative absorbance at 280 nm, 460 nm and 550 nm is shown in the chromatogram (left panel). The peak fraction (indicated by a black bar) was analyzed by SDS-PAGE (right panel). **b**, HeimdallR1 crystals in lipidic mesophase. **c**, Crystal packing of HeimdallR1. **d**, Key rhodopsin proton pump motifs in HeimdallR1. Black dashed lines indicate hydrogen-bonding interactions. Red spheres indicate water molecules. **e**–**g**, Structural comparison of HeimdallR1 with Kin4B8-XR (PDB ID: 8I2Z) and GPR (PDB ID: 7B03), from the membrane plane (**e**), the extracellular side (**f**) and the intracellular side (**g**). Notably, ICL3 of HeimdallR1 forms a membrane-extending α-helix. **h**, Interaction between ICL3 and TM6 in HeimdallR1. Black dashed lines indicate hydrogen-bonding interactions. Red spheres indicate water molecules. **i**, Sequence alignment of HeimdallRs around ICL3. The sequence alignment was created using ClustalW^74^ and ESPript 3.0^75^ servers. Boxes indicate α-helices.

**Extended Data Fig. 7.**
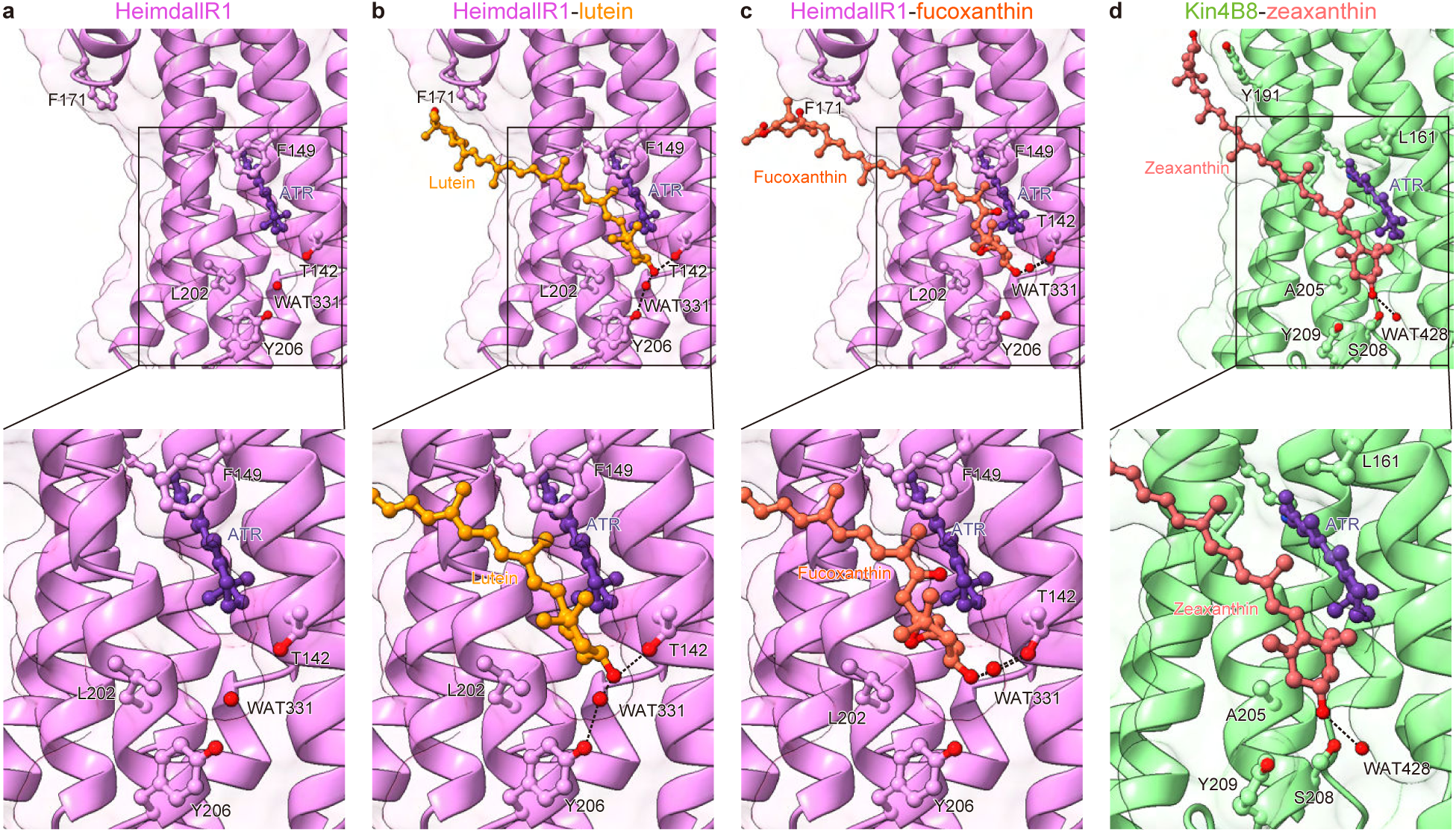
Simulation features of HeimdallR1 bound to lutein and fucoxanthin. **a,** HeimdallR1. **b,** HeimdallR1 with lutein. **c,** HeimdallR1 with fucoxanthin. **d,** Kin4B8-XR with zeaxanthin. ATR- all-*trans* retinal. Black dashed lines indicate hydrogen bonds.

**Extended Data Fig. 8.**
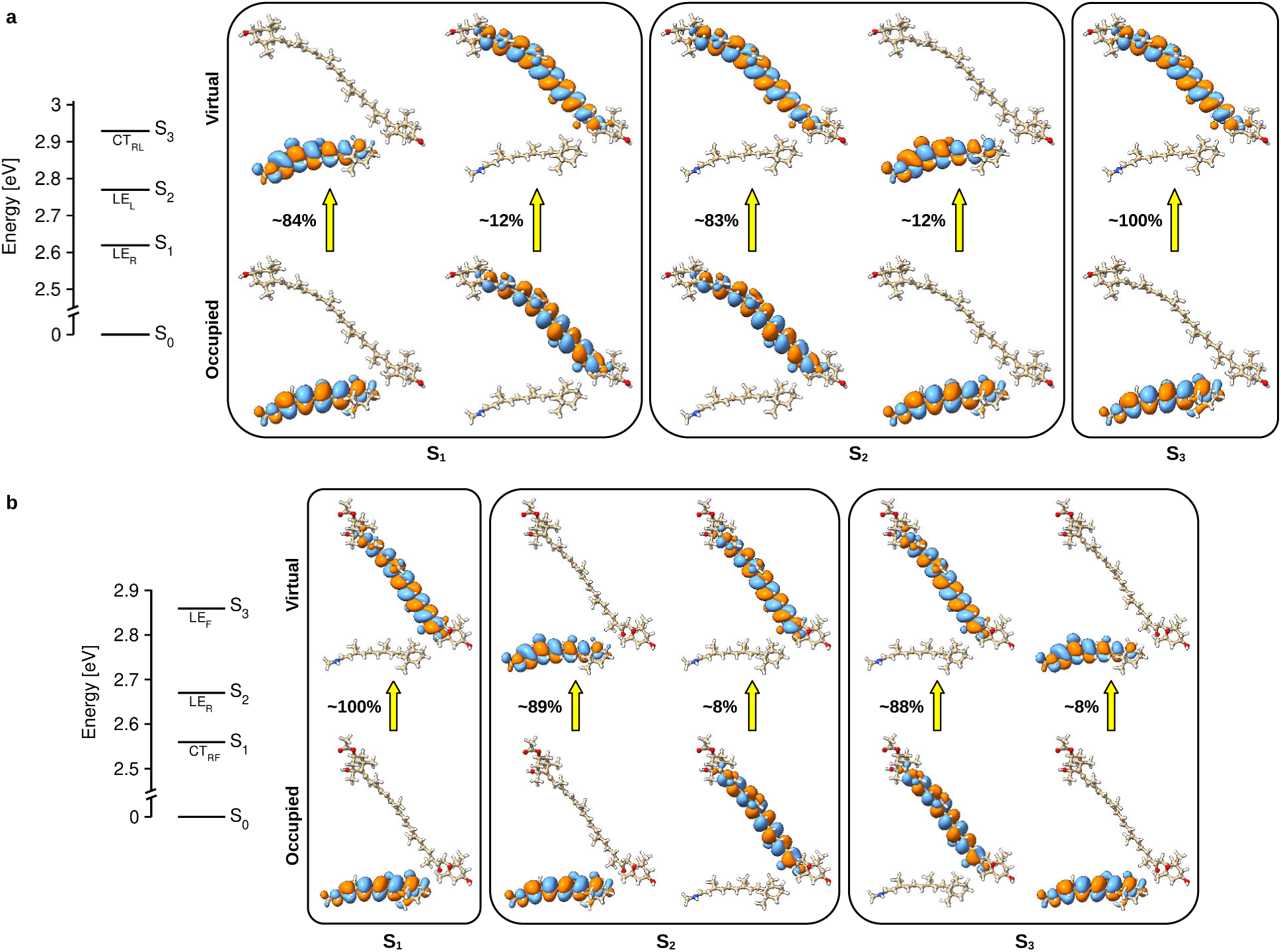
Electronic transitions in HeimdallR1/xanthophyll complexes. Energy level diagram along with the nature of the excited states (LE: local excitation; CT: charge transfer), and natural transition orbitals (isovalue=0.01 a.u.) displaying the dominant orbitals involved in the electronic transitions of the first three excited states in HeimdallR1/xanthophyll with lutein (**a**) and fucoxanthin (**b**). The subscript denotes excitation in retinal (R), lutein (L), fucoxanthin (F) or transition from retinal to the respective xanthophyll. The transition orbitals are shown only for those with contribution greater than 5% towards the excited state (transition contribution between each pair of orbitals is marked in the figure).

**Extended Data Fig. 9.**
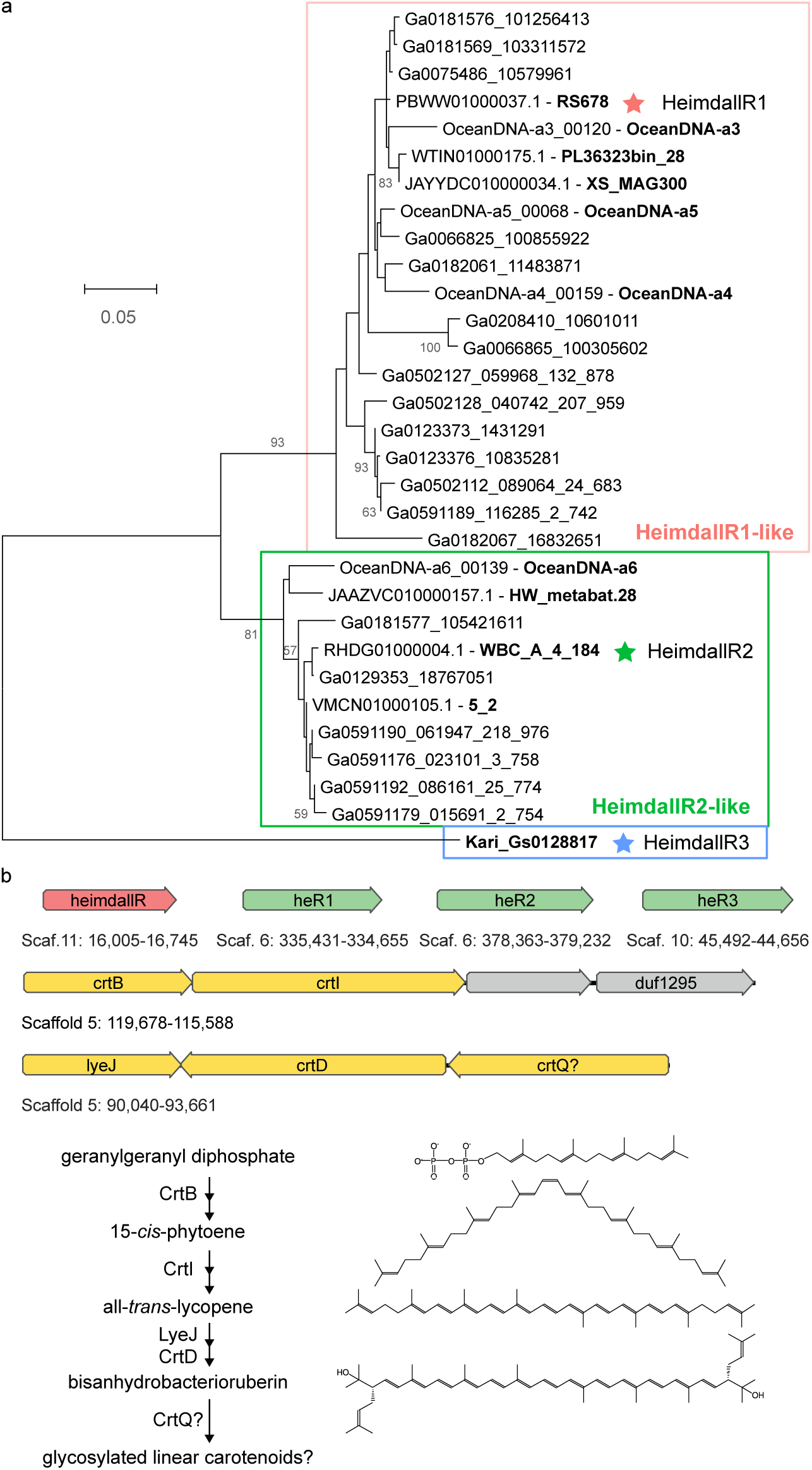
Rhodopsin genes and genes involved in biosynthesis of carotenoids in “*Ca.* Kariarchaeum”. **a**, Neighbor-joining tree of (near-)complete HeimdallR amino-acid sequences. The three clades are highlighted with color and proteins used for expression are indicated with asterisks. For proteins coming from MAGs, the corresponding scaffold names are provided. Numbers next to edges refer to bootstrap support values >50. **b**, Rhodopsins and enzymes with putative roles in carotenoid biosynthesis in the “*Ca.* Kariarchaeum pelagium” core pangenome. Top: genomic locations of the rhodopsin genes, heimdallarchaeial rhodopsins (*heimdallR*) and heliorhodopsin (*heR1-3*) genes. Middle: genomic locations of genes involved in carotenoid biosynthesis – phytoene synthase (*crtB*), phytoene desaturase (*crtI*) and two additional genes of unknown function located in the same putative operon; a putative lycopene elongase (*lyeJ,* prenyltransferase cl00337), carotenoid desaturase (*crtD,* phytoene dehydrogenase-related protein COG1233), and a putative carotenoid glycosyltransferase (*crtQ*?). Coordinates are provided for the corresponding genomic regions with respect to the pangenome scaffolds. Bottome: reactions that might be catalyzed by the corresponding enzymes.

**Extended Data Fig. 10.**
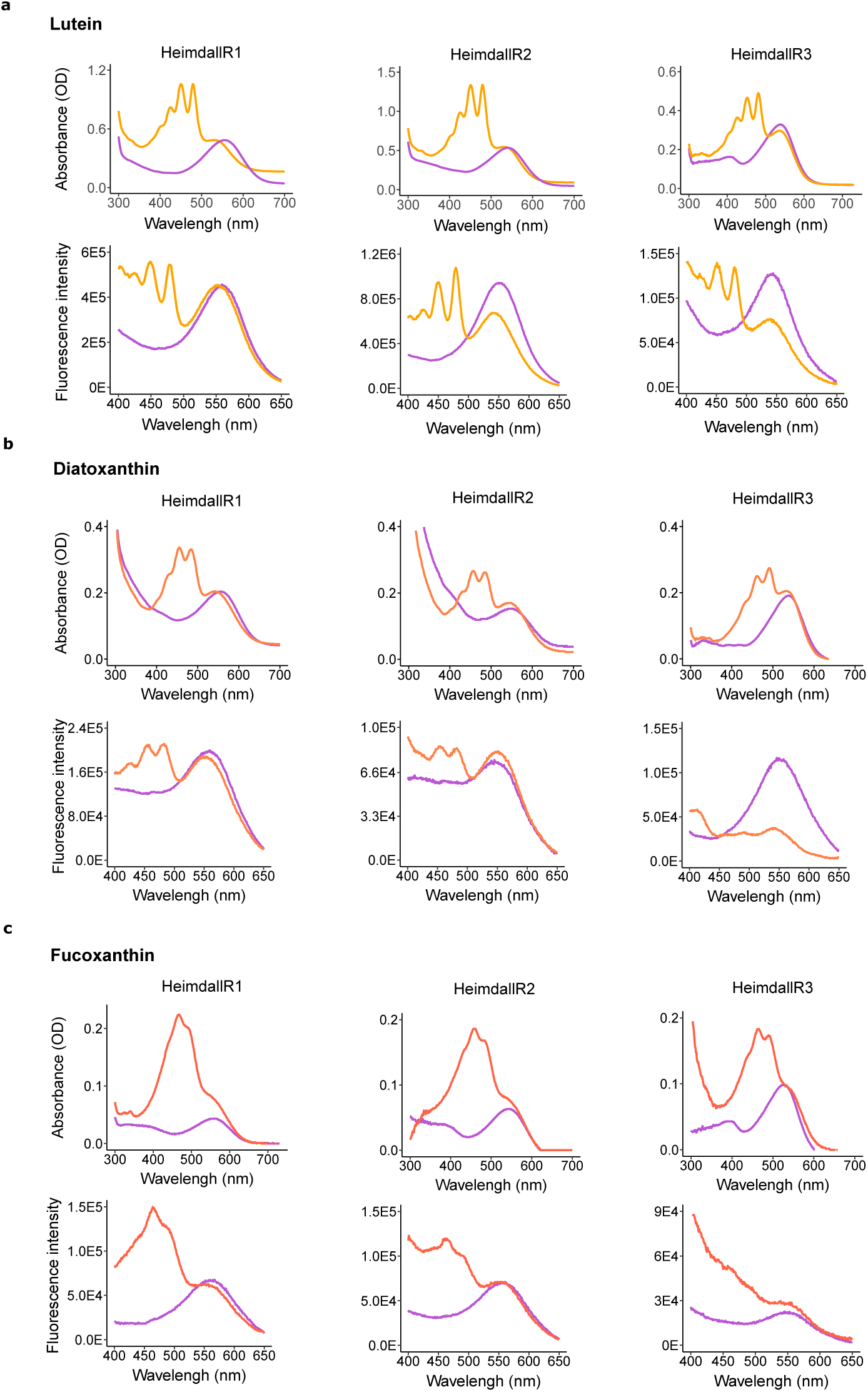
Biophysical characterization of diverse marine HeimdallRs with lutein, diatoxanthin or fucoxanthin. Absorbance change (up) and fluorescence excitation spectra (bottom) of HeimdallR1, 2 and 3 upon incubation with (orange) or without (purple) lutein (**a**), diatoxanthin (**b**), and fucoxanthin (**c**); emissions were recorded at 720 nm. The results for HeimdallR1 (already shown in Fig. 2) are presented to serve as a reference to HeimdallR2 and HeimdallR3.

## Supplementary information

Supplementary Figs. 1-3, Tables 1-3.

**Supplementary Fig. 1.**
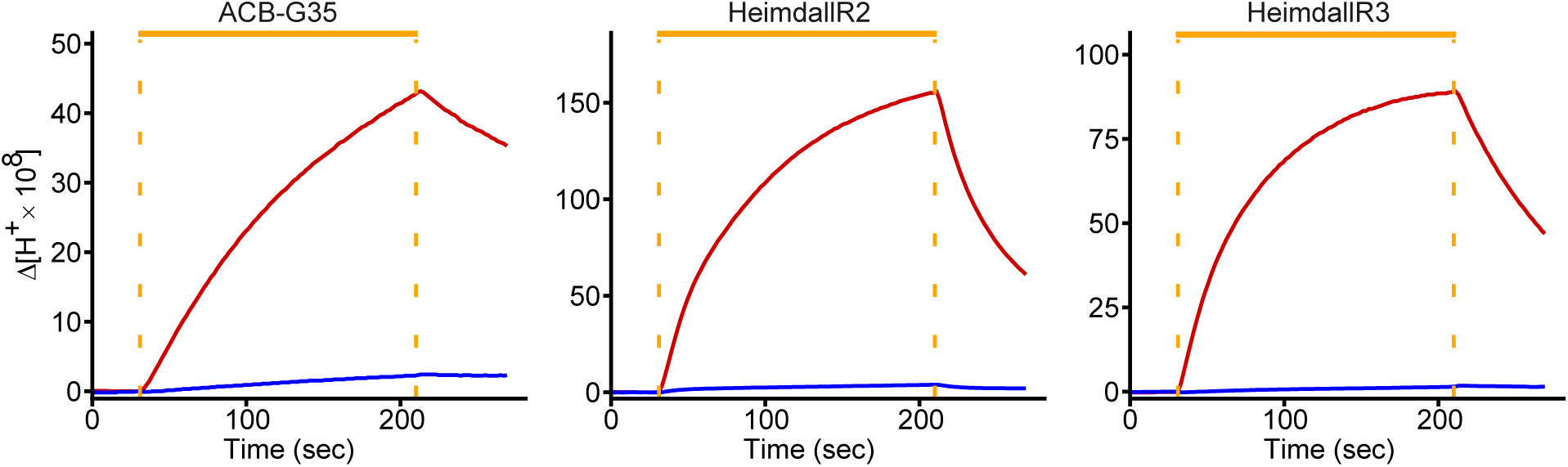
Light-driven outward proton pumping by ACB-G35 rhodopsin, HeimdallR2 and HeimdallR3. Monitoring of pH changes in *E. coli* suspension expressing ACB-G35 rhodopsin, HeimdallR2 and HeimdallR3, respectively. The rhodopsin expressing *E. coli* were illuminated with white-light for 3 min (indicated by the orange bars). Measurements were repeated under the same conditions without (red) or with (blue) the addition of 10 μM of the protonophore carbonyl cyanide-*m*-chlorophenylhydrazone (CCCP).

**Supplementary Fig. 2.**
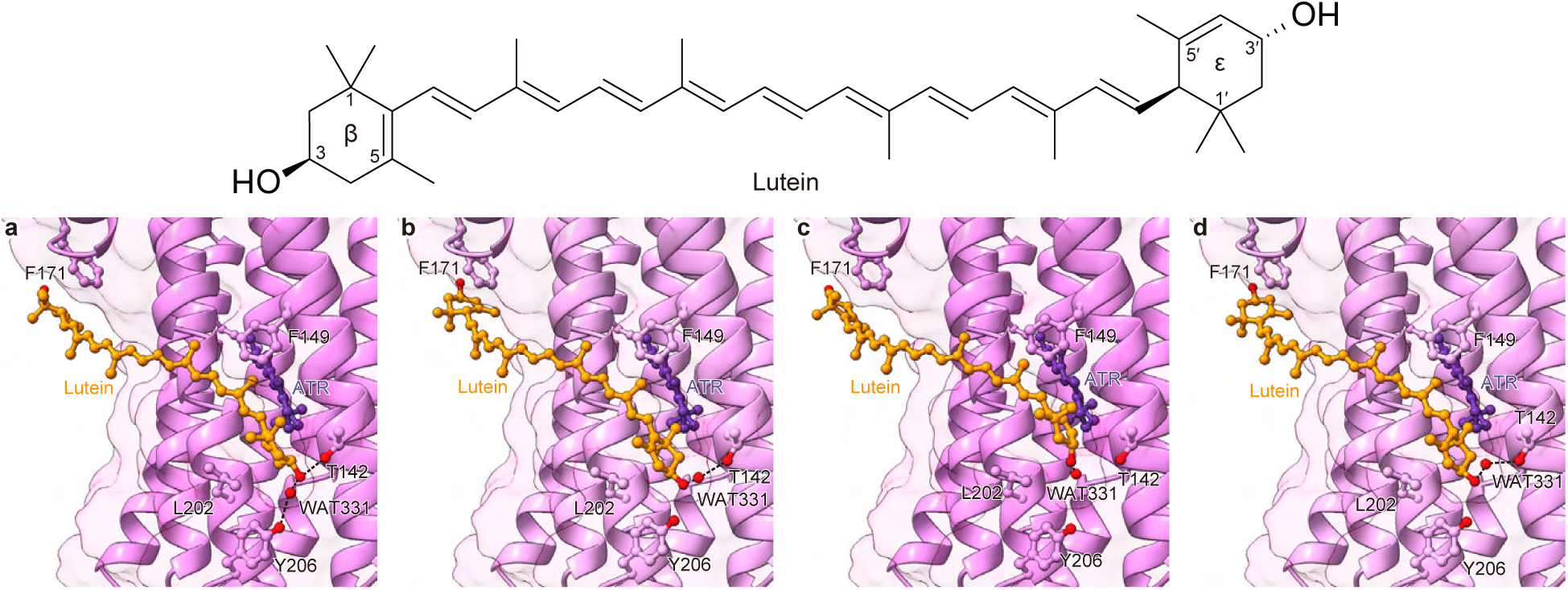
Different binding modes of lutein with HeimdallR1. Binding of lutein through the β-ionone ring with C5 pointing to the retinal (**a**) or C1 pointing to the retinal (**b**). Binding of lutein through the ε-ring with C5’ pointing towards retinal (**c**) or C1’ pointing towards retinal (**d**). Lutein structure with the β and ε rings marked is shown.

**Supplementary Fig. 3.**
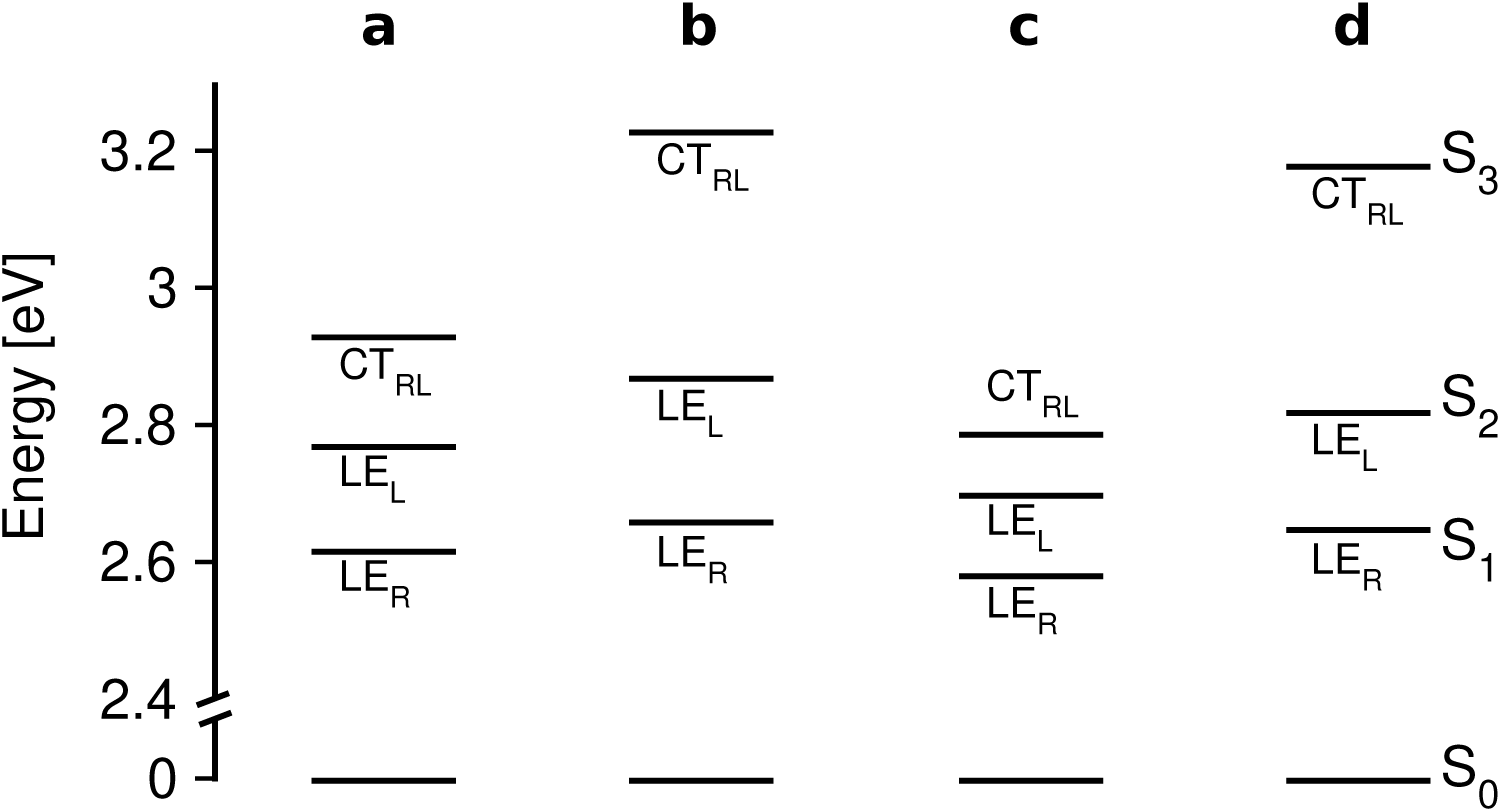
Excitation energies for different binding modes of lutein with HeimdallR1. Binding of lutein through the β-ionone ring with C5 pointing to the retinal (**a**) or C1 pointing to the retinal (**b**). Binding of lutein through the ε-ring with C5’ pointing towards retinal (**c**) or C1’ pointing towards retinal (**d**). See lutein structure in Supplementary Fig. 2.

**Supplementary Table 1.** Summary of the statistical analysis of proton pumping in HeimdallR1 expressed as log_2_-fold change (LFC, with antenna over without antenna) as a function of the presence of fenestration, antenna type and light intensity. Results of analyses of deviance (type II Wald’s χ² test) and the corresponding *p-*values are provided. FUCO — fucoxanthin, LUT — lutein, G>F — HeimdallR1-G141F mutant (fenestration blocked). Asterisks indicate significant effects. Number of measurements (ratios) used in the two designs is provided.

| Predictor | Coefficient ± standard error | Degrees of freedom | Wald's $\chi^2$ | p-value |
| --- | --- | --- | --- | --- |
| Antenna type (at 860 $\mu\text{mol m}^{-2} \text{s}^{-1}$, $n = 26$): $LFC \sim \text{antenna type} * \text{fenestration}$ | | | | |
| (intercept) | -0.0293030±0.0981457 | | | |
| antenna | 0.1624±0.1317 (FUCO) | 2 | 3.2629 | 0.1956 |
| | 0.1773±0.1317 (LUT) | | | |
| fenestration | -0.2528±0.1388 (G>F) | 1 | 12.9844 | 3.141e-04*** |
| antenna:fenestration | -0.1301±0.1913 (FUCO:G>F) | 2 | 0.2349 | 0.8892 |
| | -0.0004±0.1913 (LUT:G>F) | | | |
| Light intensity (lutein, $n = 37$): $LFC \sim \text{intensity} * \text{fenestration} + (1\vert \text{batch})$ | | | | |
| (intercept) | 0.2043±0.1125 | | | |
| intensity | -0.0001±0.0001 | 1 | 1.2159 | 0.2702 |
| fenestration | -0.5118±0.1652 (G>F) | 1 | 17.1383 | 3.475e-05*** |
| intensity:fenestration | 0.0001±0.0001 (G>F) | 1 | 0.9999 | 0.3173 |

**Supplementary Table 2.**
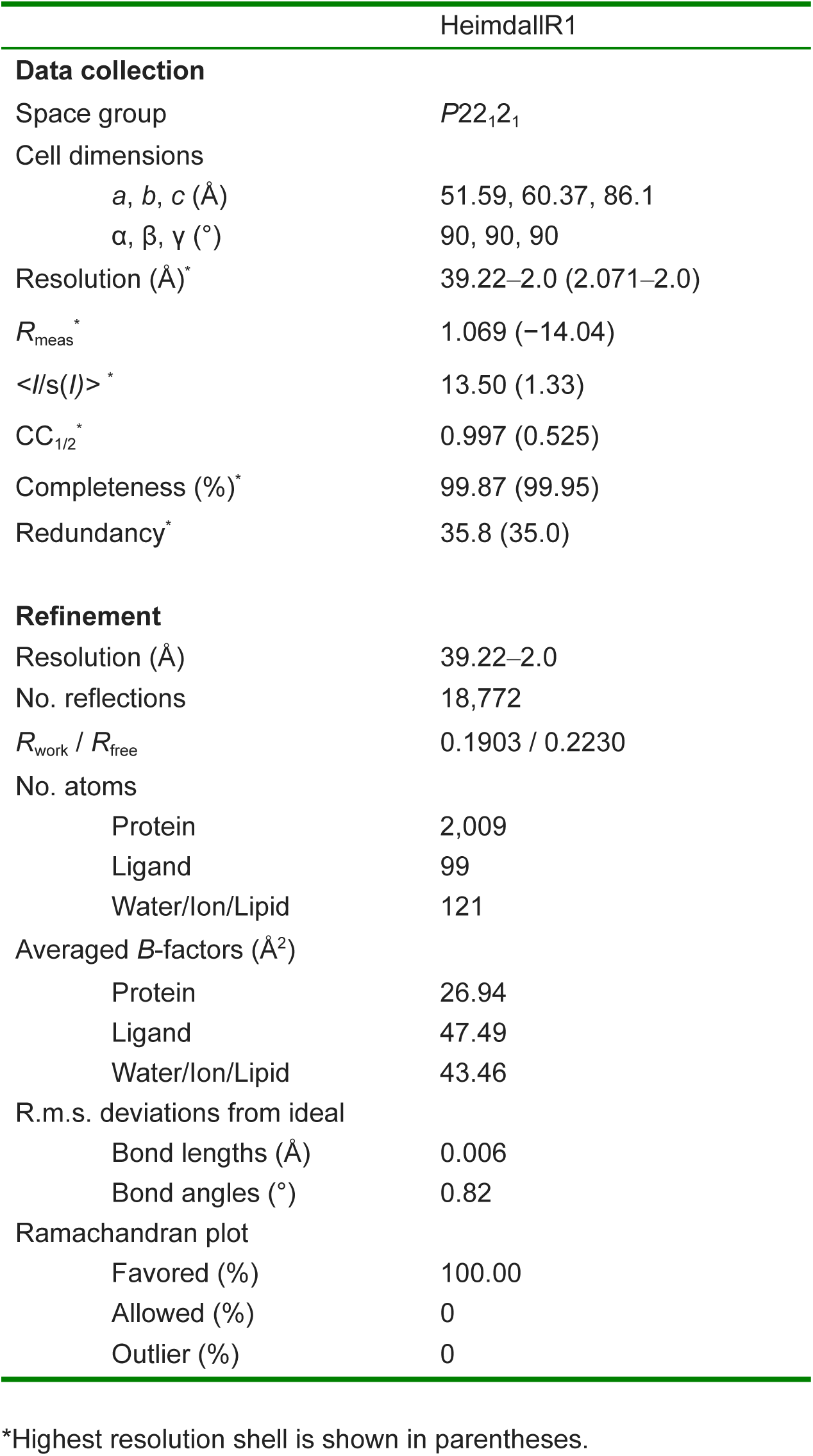
Data collection and refinement statistics.

**Supplementary Table 3.** Excitation energies (and the corresponding oscillator strengths) computed at the RI-ADC(2)/cc-pVDZ level of theory.

| State | Energy in eV (nm) | Oscillator strength |
| --- | --- | --- |
| **HeimdallR1** |  |  |
| S<sub>1</sub> | 2.43 (511) | 1.78 |
| **HeimdallR1-lutein (β-ring with C5 towards retinal)** |  |  |
| S<sub>1</sub> | 2.62 (474) | 1.11 |
| S<sub>2</sub> | 2.77 (448) | 4.20 |
| S<sub>3</sub> | 2.93 (423) | 0.00 |
| **HeimdallR1-fucoxanthin** |  |  |
| S<sub>1</sub> | 2.56 (484) | 0.00 |
| S<sub>2</sub> | 2.67 (465) | 1.38 |
| S<sub>3</sub> | 2.86 (434) | 4.32 |

